# Predicting three-dimensional genome organization with chromatin states

**DOI:** 10.1101/282095

**Authors:** Yifeng Qi, Bin Zhang

## Abstract

We introduce a computational model to simulate chromatin structure and dynamics. Starting from one-dimensional genomics and epigenomics data that are available for hundreds of cell types, this model enables *de novo* prediction of chromatin structures at five-kilo-base resolution. Simulated chromatin structures recapitulate known features of genome organization, including the formation of chromatin loops, topologically associating domains (TADs) and compartments, and are in quantitative agreement with chromosome conformation capture experiments and super-resolution microscopy measurements. Detailed characterization of the predicted structural ensemble reveals the dynamical flexibility of chromatin loops and the presence of cross-talk among neighboring TADs. Analysis of the model’s energy function uncovers distinct mechanisms for chromatin folding at various length scales.

## INTRODUCTION

The human genome contains about 2 meters of DNA that is packaged as chromatin inside a nucleus of only 10 micrometers in diameter ^2^. The way in which chromatin is organized in the three-dimensional space, i.e., the chromatin structure, has been shown to play important roles for all DNA-templated processes, including gene transcription, gene regulation, DNA replication, etc. ^3-5^. A detailed characterization of chromatin structure and the physical principles that lead to its establishment will thus greatly improve our understanding of these molecular processes.

The importance of chromatin organization has inspired the development of a variety of experimental techniques for its characterization. For example, using a combination of nuclear proximity ligation and high-throughput sequencing, chromosome conformation capture and related methods quantify the interaction frequency in three-dimensional space between pairs of genomic loci ^6, 7^, and have revealed many conserved features of chromatin organization ^1, 8-11^. A consistent picture that is emerging from these experiments is the formation of chromatin loops and topologically associating domains (TADs) at the intermediate scale of kilobases to megabases, and the compartmentalization of chromatin domains that are millions of base pairs apart in sequence. Many of the findings from these cross-linking experiments are now being validated and confirmed with microscopy imaging studies that directly probe spatial contacts ^12-19^.

Correlating one-dimensional genomics and epigenomics data with 3D contacts has been rather informative and has led to many proposals on the molecular mechanism of chromatin folding ^4, 20-24^. For example, CCCTC-binding factor (CTCF) and cohesin binding sites are frequently found at the boundaries of chromatin loops and TADs ^1, 8, 9, 11, 25^. The extrusion model takes into account of these observations to propose that cohesin molecules function as active enzymes to inch along the DNA and fold the chromatin until encountering bound CTCF ^26-28^. Epigenetic modifications are known to contribute to chromatin organization as well ^23, 29-34^. Since polymer molecules that differ in chemical compositions are known not to intermix ^35^, the distinct set of histone marks in neighboring in sequence TADs may drive the formation of chromatin domains via a phase separation mechanism ^36-38^. Comparison between these mechanistic models and experimental data has been mostly qualitative, however. Quantitative and predictive modeling of chromatin structures is in need to critically evaluate model hypotheses and to further advance our understanding of chromatin folding.

In this paper, we report the development of a predictive and transferable model to simulate the structure and dynamics of chromosomes. This model provides a high-resolution structural characterization of chromatin loops, TADs, and compartments at five-kilo-base resolution, and succeeds in quantitatively reproducing contact probabilities and power-law scaling of 3D contacts as measured in chromosome conformation capture and super-resolution imaging experiments. The transferability of this model makes it possible for *de novo* prediction of the structural ensemble for any given chromatin segment using only one-dimensional sequencing data that is available for hundreds of cell types. Detailed analysis of the model parameters and predicted structures reveals the physical principles and driving force of chromatin folding at various length scales.

## RESULTS

### Predictive modeling of chromatin organization

We introduce a predictive model to study cell-type specific 3D chromatin folding. This model takes a sequence of chromatin states derived from genome-wide histone modification profiles and a list of CTCF binding sites as input. We selected these genomic features due to their known roles in organizing the chromatin at various length scales (Fig. 1A). At the core of this model is an energy function—a force field—that is sequence specific and ranks the stability of different chromatin conformations. Starting from the input for a given chromatin segment, we use molecular dynamics simulations to explore chromatin conformations dictated by the energy function and to predict an ensemble of high-resolution structures. These structures can be compared directly with super-resolution imaging experiments or converted into contact probability maps for validation against genome-wide chromosome conformation capture (Hi-C) experiments.

**Figure 1.**
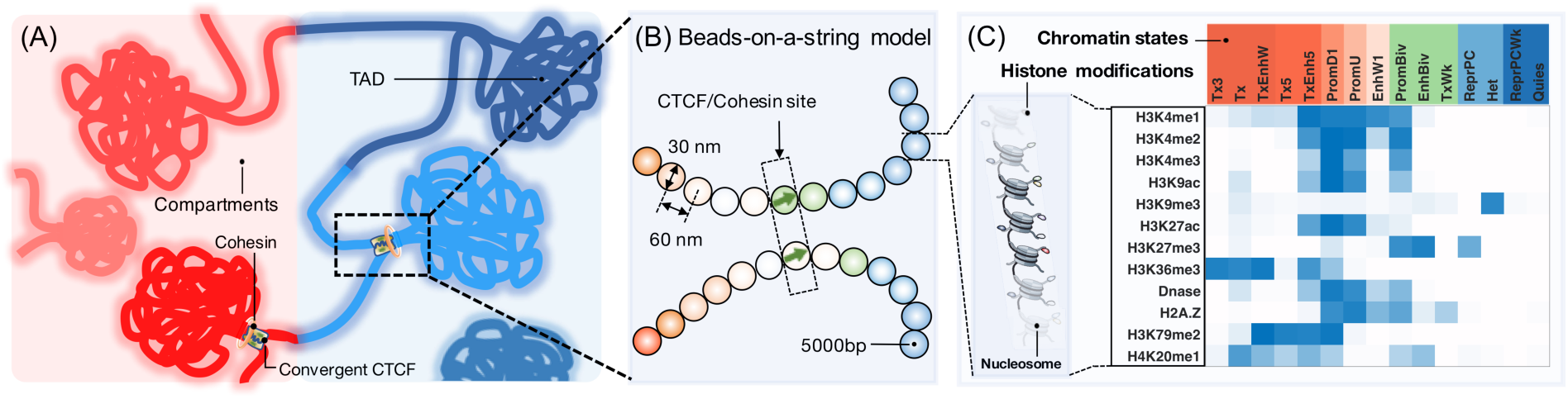
Overview of the key elements of the computational model. (A) Illustration of genome organization at various length scales that includes the formation of CTCF mediated chromatin loops, TADs, and compartments. (B) A schematic representation of the computational model that highlights the assignment of chromatin states and CTCF binding sites. Chromatin states for each bead—a 5 kb long genomic segment—are derived from the combinatorial patterns of histone marks. They are shown in part (C) as a heat map with darker colors indicating higher probabilities of observing various marks.

As shown in Fig. 1B, a continuous genomic segment is represented as beads on a string in this model. Each bead accounts for five-kilo bases in sequence length and is assigned with a chromatin state derived from the underlying combinatorial patterns of 12 key histone marks. We define a total of 15 chromatin states, identified using a hidden Markov model ^39^, to distinguish promoters, enhancers, heterochromatin, quiescent chromatin, etc. Detailed histone modification patterns for these chromatin states are shown in Fig. 1C. We further label a bead as a CTCF site if a strong binding signal is present in that location as detected by ChIP-Seq experiments (see **Methods**).

The potential energy for a given chromatin configuration ***r*** is a sum of three components, and *U*_Chrom_(***r***) = *U*(***r***) + *U*_CS_(***r***) + *U*_CTCF_(***r***). *U*(***r***) is a generic polymer potential that is included to ensure the continuity of the chromatin, and to enforce excluded volume effect among genomic loci. *U*_CS_(***r***) is a key element of the chromatin model, and is crucial to capture the formation of TADs and compartments. It quantifies the chromatin state specific interaction energies between pairs of loci. As detailed in *Section: Physical principles of chromatin organization* and **Methods**, we used a general form for *U*_CS_(***r***) to capture its dependence on genomic separation. *U*_CTCF_(***r***) is inspired by the loop extrusion model ^26-28^, and facilitates the formation of loop domains enclosed by pairs of CTCF binding sites in convergent orientation (Fig. 1A). Both *U*_CS_(***r***) and *U*_CTCF_(***r***) contain adjustable parameters that can be derived from Hi-C data following the optimization procedure developed by one of the authors ^40, 41^. Segments of chromosomes 1, 10, 19 and 21 from GM12878 cells were used for parameterization to ensure a sufficient coverage of all chromatin states (see Fig. S1). Detailed expression for the potential energy, and the parameterization procedure are provided in **Methods**.

Using the parameterized energy function, we simulated the ensemble of chromatin structures and determined the corresponding contact probability map for a 20 Mb region of chromosome 1 from GM12878 cells. As shown in Fig. 2A, the simulated contact map is in excellent agreement with the one measured by Hi-C experiments from Ref. ^1^. The Pearson correlation coefficient between simulated and experimental map exceeds 0.95. Similar comparisons for other chromosomes are provided in Fig. S2. It is evident that from these results that the simulated contact maps reproduce the overall block-wise checkboard pattern that corresponds to the compartmentalization of chromatin domains. A zoomed-in view along the diagonal of the contact map provided in Fig. 2B further suggests that chromatin loops and TADs are reproduced to a remarkable degree as well.

**Figure 2.**
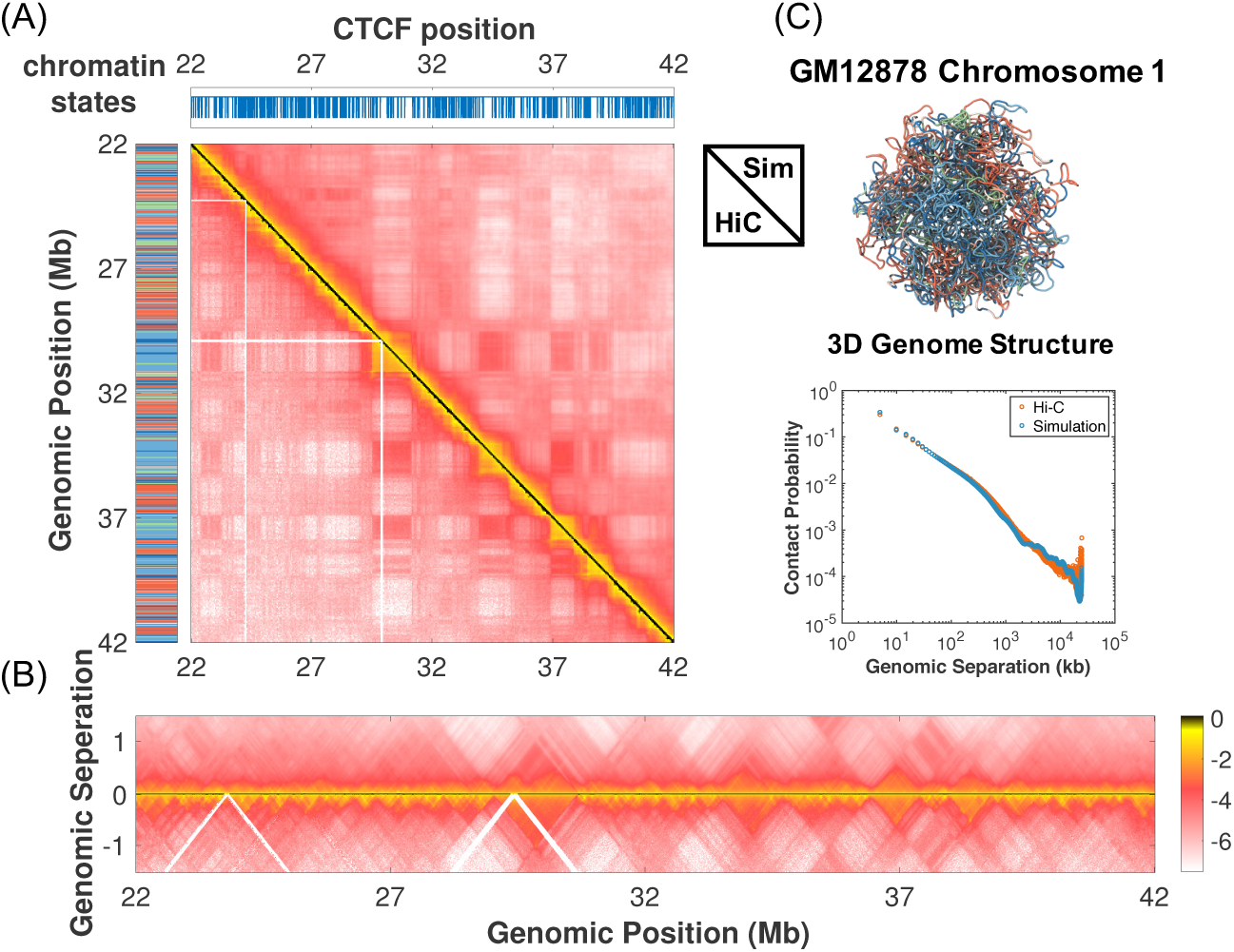
Comparison between simulated and experimental contact probability maps for a 20 Mb segment of chromosome 1 from GM12878 cells. (A) Results from simulation and the Hi-C experiment performed in Ref. ^1^ are shown in the upper and lower triangle respectively on a log scale. Also shown on the left and top panels are the sequence of chromatin states and the genomic positions of CTCF binding sites. A zoomed-in view of the contact maps along the diagonal region to highlight the formation of chromatin loops is shown in part (B). (C) A representative chromatin structure predicted by the computational model is drawn in a tube representation and colored by chromatin states. The average contact probability as a function of the genomic separation is shown below on a log-log scale for the simulated (blue) and experimental (red) contact maps respectively. The coloring scheme for chromatin states in parts (A) and (C) is consistent with that introduced in Fig. 1(C).

To more quantitatively assess the agreement between simulation and experiment, we examined multiple metrics that emphasize different features of the contact map. First, for pairs of convergent CTCF sites that are predicted by Hi-C contact maps to form loops ^1^, we calculated the contact enhancements as the ratio of their contact probabilities over the mean contacts averaged over a locally selected background region (see *SI Section: Contact enhancement metric for chromatin loops* for detail*s*). As shown in Fig. S3, the contact enhancements are larger than one for nearly all (over 92%) of the pairs, which indicates a strong enhancement of spatial colocalization between loop anchors. In the meantime, for randomly selected genomic pairs with comparable sequence separations as those found in chromatin loops, only approximately 50% of them exhibit contact enhancements larger than one. Furthermore, over 74% of the loop pairs exhibit a contact enhancement that is larger than the 90th percentile of the random distribution.

We next determined the correlation coefficients between the top five eigenvectors for simulated and experimental contact matrices. As shown in Fig. S4, the contact maps reconstructed using only these eigenvectors recapitulate the formation of TADs and compartments observed in the original maps. The high correlation between simulated and experimental eigenvectors (with Pearson correlation coefficients at approximately 0.8) supports that the corresponding features are well captured by the computational model, and confirms the qualitative observations from Fig. 2.

To demonstrate the transferability of the computational model across chromosomes and cell types, we performed additional simulations for chromosomes from GM12878, K562, and Hela cells, whose Hi-C data were not included during the parameterization procedure. As shown in Fig. 3 and Fig. S5, these *de novo* predictions are again in good agreement with experimental results as measured by Pearson correlation coefficients and other quantitative metrics. We emphasize that the sole inputs for these simulations are chromatin states derived from histone modification patterns and genomic positions and orientations of CTCF binding sites. The excellent performance of these *de novo* predictions demonstrates that the chromatin model introduced here provides a consistent description of the 3D genome organization across cell types.

**Figure 3.**
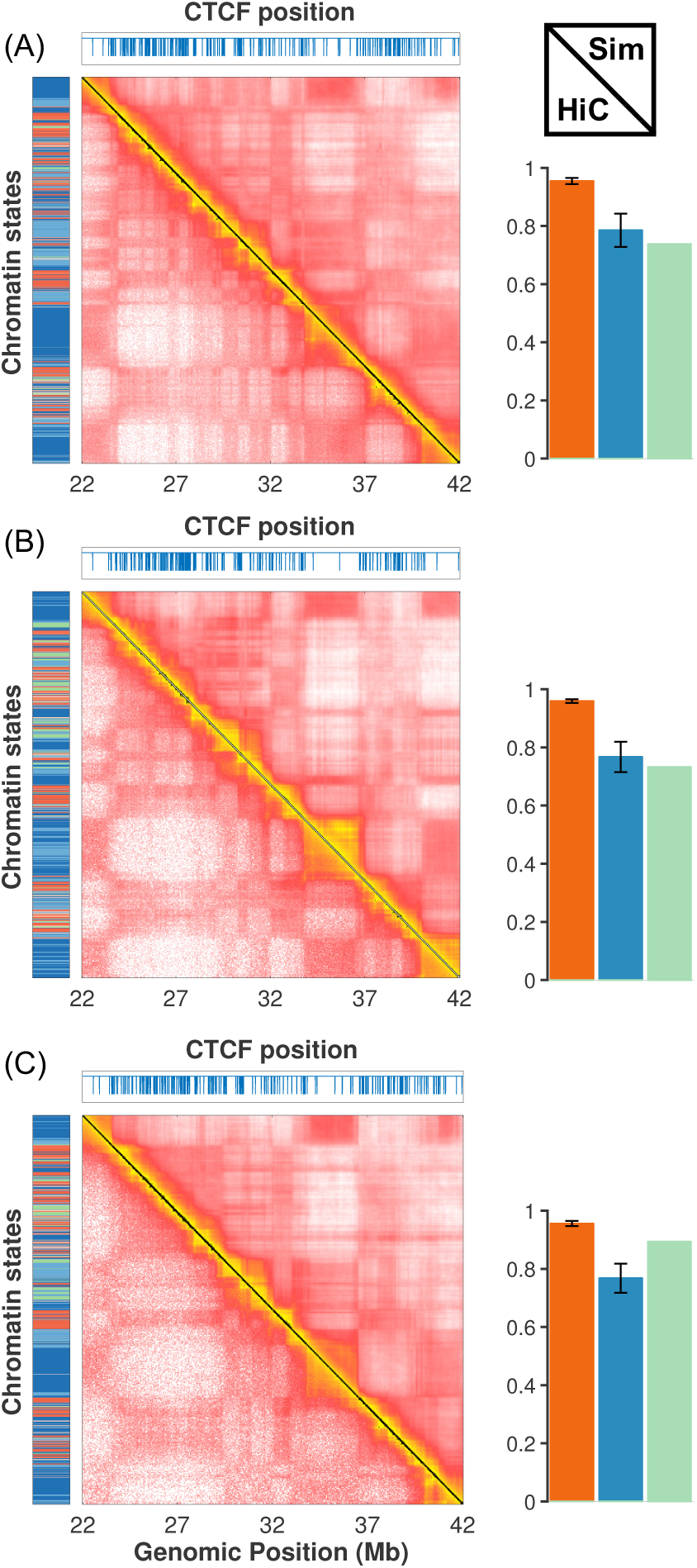
Transferability of the computational model across chromosomes and cell types. (*Left*) Comparison between simulated and experimental contact maps for chromosome 2 from GM12878 (A), K562 (B), and Hela cells (C). (*Right*) Quality of computational predictions for all chromosomes from the three cell types measured by quantitative metrics that include average Pearson correlation coefficients between simulated and experimental contact maps (red), average correlation coefficients between the top five eigenvectors for the logarithm of contact matrices (blue), and percent of chromatin loops with contact enhancements larger than the 90th percentile of the contact enhancement distribution for randomly selected genomic pairs (green).

### Structural characterization of chromatin organization

We next analyze the simulated 3D structural ensembles to gain additional insights on chromatin organization. Consistent with previous experimental and theoretical studies ^36, 42, 43^, our model reproduces the clustering of active genes and their preferred location at the exterior of the chromosome. This clustering is evident from the representative structure shown in Fig. 2, and from the quantitative analysis over the entire structure ensemble provided in Fig. S6.

Super-resolution imaging experiments probe chromatin organization in 3D space to quantify spatial distances between genomic segments. These 3D measurements can be compared directly with simulated chromatin structures, and thus provide a crucial validation of the computational model parameterized from Hi-C experiments with independent datasets. To understand the overall compactness of various chromatin types, we selected a set of active, repressive and inactive chromatins and determined their radiuses of gyration from the ensemble of simulated structures. These different chromatin types are identified using two key histone marks H3K4me2 and H3K27me3 (Fig. 4A). The complete list of chromatin domains with their genomic locations is provided in the Extended Data Sheet. As shown in Fig. 4B, the radius of gyration increases at larger genomic separation following a power law behavior in all cases with exponents of 0.34, 0.31 and 0.23 for the three chromatin types respectively. These scaling exponents are in quantitative agreement with imaging measurements performed for Drosophila chromosomes ^44^ and support the notion that active chromatins adopt less condensed conformations to promote gene activity. Consistent with the imaging study performed on chromosome 21 from IMR90 cells ^13^, we also observe a strong correlation between Hi-C contact probabilities and spatial distances for pairs of genomic loci (Fig. 4C).

**Figure 4.**
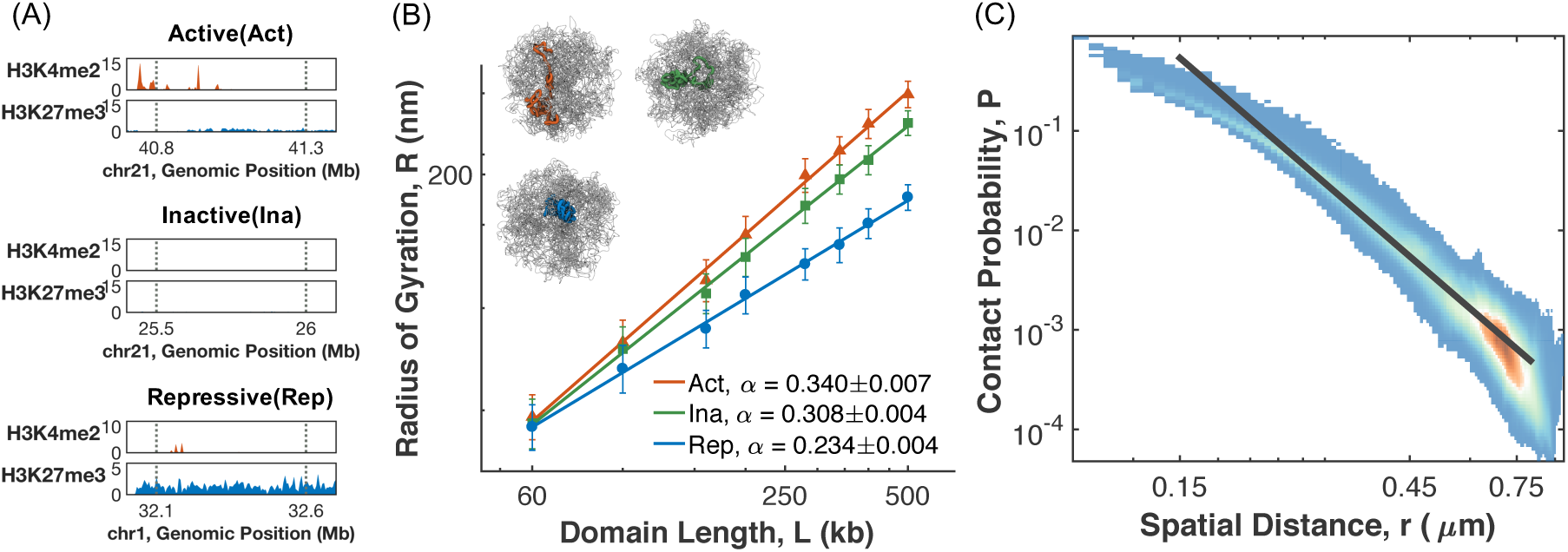
Simulated chromatin structures reproduce findings from super-resolution microscopy experiments. (A) Characteristic histone modification profiles for repressive, active and inactive chromatin. (B) The sizes of repressive (blue), active (orange) and inactive (green) chromatin domains, as measured by their radiuses of gyration, are plotted as a function of the genomic separation on a log scale. The straight lines correspond to numerical fits of the data with a power-law expression *R* = *R*_*o*_ *L*^*α*^, with the values of *α* shown in the legend. Representative structures of 500 kb in length for the three chromatin types are shown in the inset. (C) Scatter plot of the contact probabilities between pairs of genomic loci versus their spatial distances shown on a log-log scale. The black line is the best fit to the data using the expression *P* = *P*_*o*_*r*^*β*^, with *β* = −4.18.

One of the most striking features revealed by high-resolution Hi-C experiments is the formation of chromatin loops anchored at pairs of convergent CTCF sites ^1, 9, 45, 46^. Microscopy studies that directly visualizes 3D distances using fluorescence in situ hybridization (FISH) methods further find that these loops are dynamic, and despite their high contact frequencies, loop anchors are not in close contact in every cell ^16, 47-49^. Consistent with their dynamic nature, chromatin loops in our simulation adopt flexible conformations as well. As shown in Fig. 5A, for the loop formed between chr1:37.76-37.93 Mb, we observe a large variance in the probability distribution of its end-to-end distances. Two example configurations of the loop domain with distance at 0.08 and 0.24 *μm* are shown in the inset. A systematic characterization of all the loops identified in Ref. ^1^ for the simulated chromatin segment shows that the conformational flexibility is indeed general, though there is a trend in decreasing variance for loops with larger contact probabilities (Fig. 5B).

**Figure 5.**
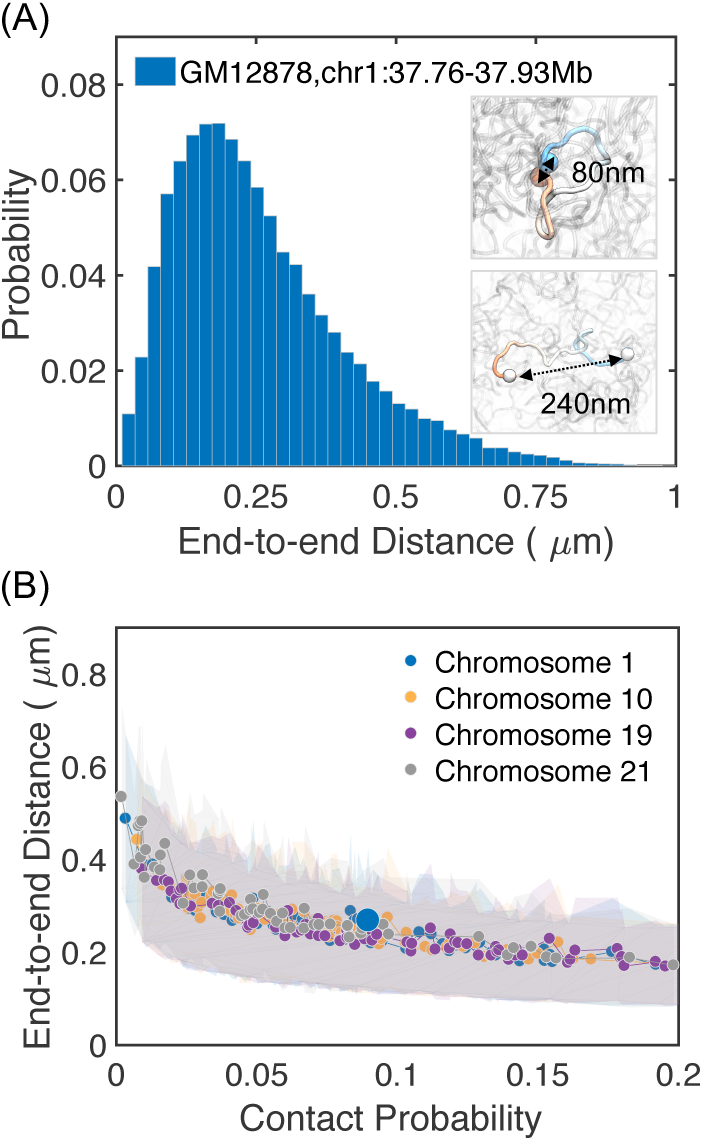
Structural characterization of chromatin loops. (A) Probability distribution of the end-to-end distance for the chromatin loop formed between chr1:31.43Mb and chr1:31.61Mb from GM12878 cells. Two example configurations that correspond to open and closed loop structures are shown in the inset. (B) End-to-end distances of chromatin loops versus their corresponding contact probabilities. The shaded areas represent the variances in distances estimated from the simulated structural ensemble.

Compared to chromatin loops, TADs are longer and are stabilized by a complex set of interactions ^24^. The analysis of their structural ensemble is less straightforward, and the end-to-end distance may not be sufficient for a faithful description of their conformational dynamics ^50^. We, therefore, applied a machine learning approach to automatically identify reaction coordinates that capture statistically significant conformational fluctuations present in the simulated structural ensemble. Borrowing ideas from protein folding studies, we approximate these reaction coordinates using collective variables with slowest relaxation timescales as determined following the diffusion map analysis ^51, 52^. Diffusion map analysis has been successfully applied to a variety of systems to provide mechanistic insights on the conformational dynamics involved in protein folding, ligand diffusion, etc. ^53, 54^.

We applied the diffusion map technique to the predicted structural ensemble of the genomic region chr1:34-38Mb from GM12878 cells that consists of three visible TADs. As shown in Fig. 6, multiple basins are observed in the probability distribution for the two identified reaction coordinates, suggesting significant conformational rearrangements. To gain physical intuition on the reaction coordinates, we calculated the corresponding contact maps at various regions. As shown in the top panel, reaction coordinate one captures the formation of contacts between TAD1 and TAD3 while the structures for all three TADs remain relatively intact. On the other hand, progression along reaction coordinate two (left panel) leads to significant overlaps between TAD1 and TAD2. Interaction between TAD2 and TAD3 can also be observed along a third coordinate as shown in Figure S7B. Example structures for the three TADs in various regions are also provided on the right panel. These results are consistent with the notion that TADs are stable structural units for genome organization ^24^, but also suggest the presence of significant cross-talk among neighboring TADs ^55^.

**Figure 6.**
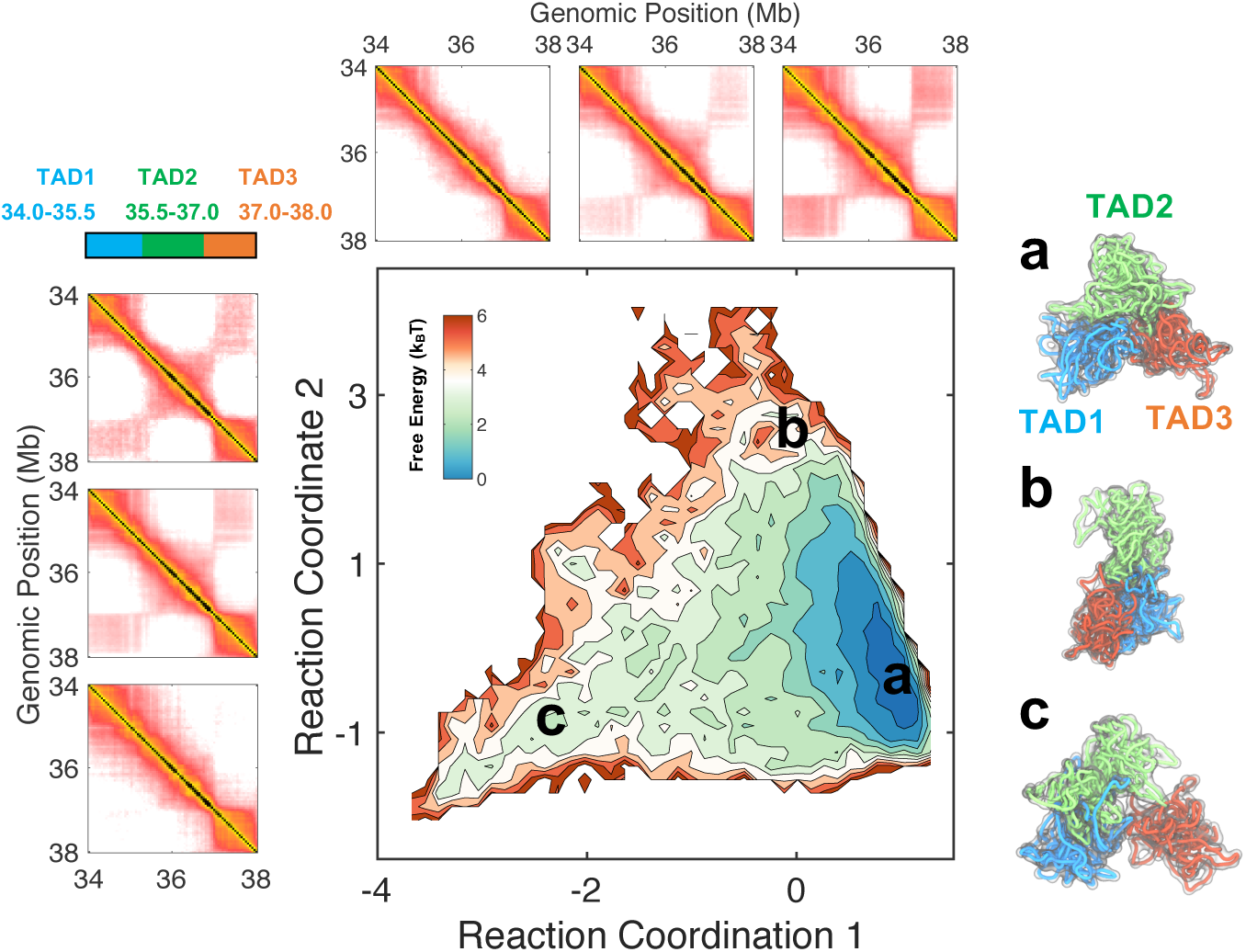
Structural characterization of topologically associating domains using the diffusion map technique. (*Center*) Free energy profile of TAD conformations projected onto two coordinates that describe the slowest collective motions. The (*Left*) and (*Top*) panels illustrate the change in contact maps along the two coordinates. (*Right*) Representative structures for the chromatin segment at various positions indicates in the central and bottom panel. The three contact maps for reaction coordinate 1 were calculated using chromatin structures that fall into the regions [−2.5, −0.5), [−0.5,0.5) and [0.5, 1.5). The three regions used to calculated the contact maps for reaction coordinate 2 are [−2.5, −1.0), [−1.0, 1.5), and, [1.5,3.5).

### Physical principles of chromatin organization

Though the exact molecular mechanism and driving force for chromatin folding remain elusive, it is becoming increasingly clear that different molecular players are involved in organizing the chromatin at various length scales ^20, 56^. For example, transcription factors and architectural proteins are critical in stabilizing the formation of chromatin loops and TADs ^4, 24, 25^. On the other hand, nuclear compartments, such as the nucleolus and the nuclear envelope, contribute to chromatin compartmentalization and mediate contacts among chromatin domains separated by tens of Mb in sequence ^21, 57^. We expect that these different molecular mechanisms will give rise to distinct interaction energies at various genomic length scales. For example, for the same pair of chromatin states, as the genomic separation between them is varied, the interaction energy that stabilizes their contact should vary. In the following, we examine the dependence of inferred contact energies on genomic separation to reveal the principles of genome organization.

Fig. 7A presents the derived contact energies among chromatin states *U*_CS_(***r***) at various genomic separations (500kb, 1.5Mb, 4Mb and 10 Mb from left to right), with blue and red for attractive and repulsive interactions respectively. A notable feature for all four length scales is the clear partition of chromatin states into at least two groups that correspond to well-known active and repressive chromatins respectively. For example, attractive interactions are observed among the top half chromatin states that include promoters (PromD1, PromU), enhancers (TxEnh5, Enhw1) and gene body (Tx), and for the bottom half that include inactive chromatin (Quies), polycomb repressed domain (ReprPC) and heterochromatin (Het). The unfavorable interactions among active and repressive chromatins will drive their phase separation shown in Fig. 2C and Fig. S2C. Partitioning of chromatin states into active and inactive groups is also evident from the eigenvectors for the largest in magnitude eigenvalue of the interaction matrices shown in Fig. 7B, with grey for positive and red for negative values.

**Figure 7.**
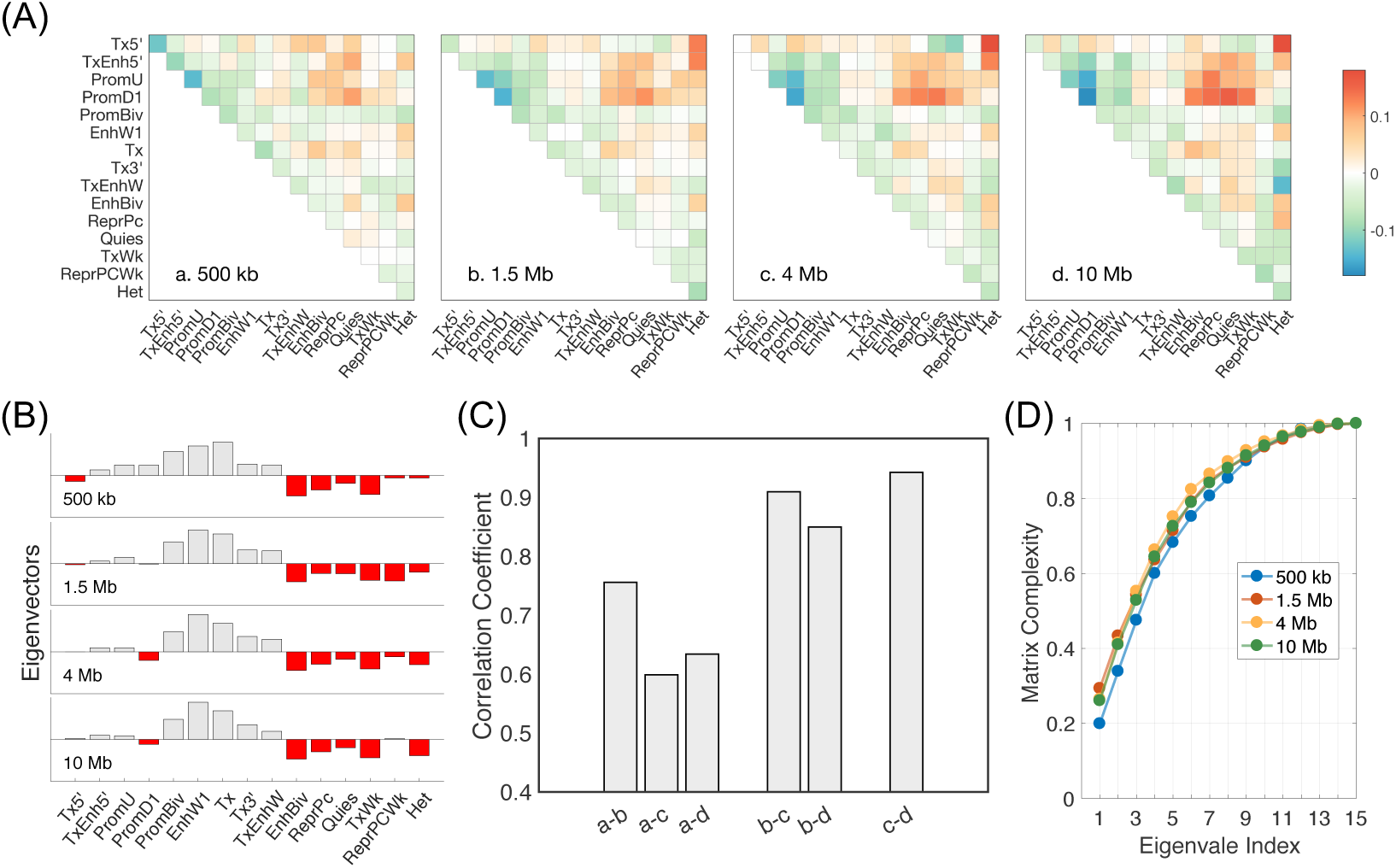
Dependence of chromatin state interaction energies on genomic separation. (A) Heat maps for the interaction matrices at various genomic separations, with blue and red corresponding to attractive and repulsive interactions respectively. We subtracted out the mean of the interaction energies in order to shift different plots to the same scale. (B) The eigenvectors corresponding to the largest eigenvalues of the four interaction matrices, with grey and red indicating positive and negative values respectively. (C) Pearson correlation coefficients between interaction matrices at different scales. (D) The complexity measure for different interaction matrices as a function of the index for top eigenvalues. See text for the definition of the complexity measure.

Despite their overall similarities, the interaction energies at various genomic separations differ from each other. To quantify their differences, we determined the pairwise Pearson correlation coefficients between the interaction matrices. As shown in Fig. 7C, the interactions that are responsible for TAD formation (∼ 1 Mb) indeed differ significantly from those that lead to chromatin compartmentalization (∼ 10 Mb), as evidenced by the low correlation among them. Strikingly, the correlation coefficient between interaction matrices at 4 Mb and 10 Mb exceeds 0.9, indicating the convergence of chromatin interactions at large genomic separation.

We further compared the complexity of the interaction matrices by calculating the ratio of the first *n* eigenvalues over the sum of all eigenvalues. Fig. 7D plots this complexity measure as a function of *n*, and absolute values of the eigenvalues were used to calculate the measure. For all three matrices with genomic separation larger than 1 Mb, we find this ratio is approximately 80% at *n* = 6. Therefore, overall features of these matrices can indeed be explained with the first six eigenvectors, explaining the success of our previous effort in modeling chromatin organization with six compartment types ^36^. However, more eigenvectors are needed, especially for short range in sequence interactions, to capture the full matrix complexity. These results together highlight the presence of distinct mechanisms that fold the chromatin at various genomic separations, and argues the importance of using sequence length dependent contact energies.

## DISCUSSION

We introduced a computational model for *de novo* prediction of chromatin structures and demonstrated its transferability across chromosomes and cell types. The model input is derived from genome-wide histone modification profiles that are available for hundreds of cell types via the epigenome roadmap project ^58^. We, therefore, anticipate a straightforward application of this model to characterize the differences of chromatin structures across cell types and to understand the role 3D genome organization in cell differentiation and cell fate establishment.

Histone modifications have long been recognized as crucial for the genome’s function^59^. The “histone code” hypothesis was proposed to rationalize the presence of numerous types of histone marks and the importance of their combinatorial roles ^60^. However, a mechanistic understanding of the relationship between these chemical modifications and the functional outcome remains lacking ^61^. The success of the computational model introduced here in predicting chromatin structures argues for the importance of histone modifications in organizing the genome. It is tantalizing to hypothesize that the histone code can be understood from a structural perspective. Epigenome engineering experiments that perturb histone modifications at specific genomic locations will be helpful to elucidate further whether the relationship between 1D histone modifications and 3D genome organization is causal.

## ACKNOWLEDGMENTS

This work was supported by National Science Foundation Grants MCB-1715859. Y.F.Q. acknowledges Lester Wolfe Fellowship for financial support.

## AUTHOR CONTRIBUTIONS

Y.F.Q. and B.Z. performed research, analyzed data and wrote the manuscript.

## METHODS

### Energy function of the chromatin model

The energy function of the chromosome model, which can be rigorously derived following the maximum entropy principle ^40, 41^, adopts the following form

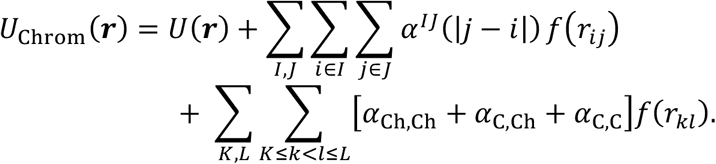

*U*(***r***) defines the generic topology of the chromosome as a confined polymer with excluded volume effect. The second term incorporates the sequence length dependent contact energies *α*^78^(|*j* − *i*|) between pairs of loci *i, j* characterized with chromatin states *I, J* respectively. As discussed in the main text, the dependence of contact energies on sequence length separation is crucial to reproduce the hierarchical genome organization, and to detect independent mechanisms of chromatin folding at different length scales. *f*(*r*_*ij*_) measures the contact probability between a pair of loci *i* and *j* separated by a distance *r*_*ij*_. Finally, the last term, inspired by the recently proposed extrusion model ^26-28^, is included to model the formation of chromatin loops. In particular, the genomic segment enclosed by a pair of convergent CTCF binding sites experiences a condensing potential due to the binding of cohesin molecules. We limit this potential to convergent CTCF pairs that are separated by no more than 4 CTCF binding sites with 5’ – 3’ orientation or 4 CTCF binding sites with 3’ – 5’ orientation to mimic the finite processivity of cohesin molecules ^27^. For generality, three different potentials are used for CTCF-CTCF interaction (*α*_C,C_), CTCF-chromatin interaction (*α*_C,Ch_) and chromatin-chromatin interaction (*α*_Ch,Ch_). Explicit mathematical expressions of the potential energy function are provided in the supporting information, and numerical values of the parameters are provided in the Extended Data Sheet.

### Simulation details

We carried out constant temperature simulations to predict chromatin structures consistent with the energy function *U*_chrom_(***r***) using the molecular dynamics software package LAMMPS ^62^. For each contact map presented in the manuscript, a total of eight independent 40-million-timestep long simulations were performed to ensure sufficient statistics. More details on the simulation are provided in the supporting information.

To enable a quantitative comparison between simulated chromatin structures with microscopy imaging data, we estimate a 5kb long genomic segment with a width of 30 nm and a length of 60 nm based on a high-resolution chromatin structure characterized by cryogenic electron microscopy (Cryo-EM) technique ^63^.

### Chromatin states from epigenomics data

A key input of the computational model is the sequence of chromatin states that captures the variation of epigenetic modifications along the genome sequence. Following Ref. ^39^, we defined chromatin states as the set of unique combinatorial patterns of histone marks. Using a multivariate hidden Markov model that maximizes the posterior probability of assigning a hidden state to each genomic segment given the sequence of observed histone modifications ^64^, we derived 15 chromatin states from genome-wide profiles of 12 key histone marks collected from six cell types. The dataset used for chromatin state inference is listed in the Extended Data Sheet. Detailed histone modification patterns for these chromatin states are shown in Fig. 1C. With the set of chromatin states specified, every five-kilo-base long segment can then be assigned to a chromatin state based on its histone modification profiles, and a sequence of chromatin states for the entire chromatin segment can be defined as the simulation input.

### Genomic locations and orientations of CTCF binding sites from ChIP-Seq data

To capture the formation of chromatin loops, we compiled a list of CTCF-binding sites along the chromatin of interest using cell-type specific ChIP-Seq data (see Extended Data Sheet). As both CTCF and cohesin molecules are found at the boundaries of the majority of chromatin loops ^9^, only CTCF binding sites that have a bound cohesin subunit Rad21 within a 50 base range were included in this list. We then determined the orientation of these binding sites by aligning them with the CTCF motifs reported in Ref. ^65^. When this alignment fails, and no CTCF motifs can be found near the binding site, we assigned CTCF binding site orientations based on the relative positions of the nearest Rad21 peaks (see the *SI Section: CTCF binding sites from ChIP-Seq data* for details).

### Diffusion map analysis

For molecular systems that exhibit a separation of timescales, it is often desirable to approximate their dynamics at long time limit with a handful of slow variables. The time evolution of these slow variables should be Markovian and independent of the fine details of the high dimensional system to capture the dynamical behavior of the system on a coarsened timescale. Mathematically it has been proven that an optimal choice of these slow variables is the first few eigenfunctions of the backward Fokker–Planck diffusion operator ^51^. Diffusion map is a data-driven approach that approximates these eigenfunctions and therefore the slow variables by defining a random walk process on the simulation data ^66^.

In particular, for *N* chromatin configurations selected from the simulated structural ensemble, we first constructed a transition probability matrix *K* for the random walk by defining its elements as

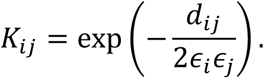

The eigenfunctions of the above transition matrix can be shown to converge to that of the Fokker–Planck operator in large *N* limit. The distance between two configurations *d*_*i*_ was calculated as the mean difference of their corresponding contact probability maps. We followed the algorithm proposed in Ref. ^52^ to normalize the matrix and to estimate *ϵ*_*i*_. From the normalized transition matrix, we then determined its eigenfunctions and used the top two with smallest non-zero eigenvalues as the reactions coordinates shown in Fig. 6 (see Fig. S7 for eigenvalues).

