## Supplementary Materials for "Predicting three-dimensional genome organization with chromatin states"

### Supporting Information for "Predicting three-dimensional genome organization with chromatin states"

**Energy function of the chromatin model.** As mentioned in the main text, we model the chromatin as beads on a string. The potential energy for a given chromatin configuration $\boldsymbol{r}$ consists of three components

$$U_{\text{Chrom}}(\boldsymbol{r}) = U(\boldsymbol{r}) + U_{\text{CS}}(\boldsymbol{r}) + U_{\text{CTCF}}(\boldsymbol{r}) \qquad [1]$$

$U(\boldsymbol{r})$ is the energy function for a confined homopolymer that consists of the following four terms, $U_{\text{bond}}$, $U_{\text{angle}}$, $U_{\text{sc}}$ and $U_{\text{c}}$.

$U_{\text{bond}}$ is the bonding potential between neighboring beads and is defined as

$$U_{\text{bond}} = K_2(r - r_o)^2 + K_3(r - r_o)^3 + K_4(r - r_o)^4,$$

where $K_2 = K_3 = K_4 = 20\frac{\epsilon}{\sigma^2}$ and $r_o = 2.0\ \sigma$. $\sigma = 30$ nm is the diameter of the bead. We choose the distance between neighboring beads as 60 nm to arrive at a nucleosome line density $0.43\ nm^{-1}$ that is consistent with the recent chromatin fiber structure resolved at 11 Å with cryo-EM [2]. $\epsilon = k_B T$ defines the energy scale of the model.

$U_{\text{angle}}$ is an angular potential that defines the persistence length of the polymer. It is applied to all connected three consecutive monomers in the following form

$$U_{\text{angle}} = K_a[1 - \cos(\theta - \pi)],$$

where $K_a = 2\ \epsilon$.

$U_{\text{sc}}$ is a soft-core potential applied to all the non-bonded pairs to enforce the excluded volume effect among genomic loci, and is defined as

$$U_{\text{sc}} = \begin{cases} 0.5E_{\text{cut}}\left(1 + \tanh\left[\frac{2U_{\text{LJ}}(r)}{E_{\text{cut}}} - 1\right]\right), & r \le r_{\text{cut}} \\ U_{\text{LJ}}(r), & r_{\text{cut}} \le r \le \sigma 2^{1/6}, \\ 0, & r > \sigma 2^{1/6}. \end{cases}$$

The above expression corresponds to the Lennard-Jones potential $U_{\text{LJ}}(r) = 4\epsilon\left[\left(\frac{\sigma}{r}\right)^{12} - \left(\frac{\sigma}{r}\right)^{6}\right] + \epsilon$ capped off at a finite volume within a repulsive core to allow for chain crossing at finite energetic cost. $r_{\text{cut}}$ is chosen as the distance at which $U_{\text{LJ}}(r) \equiv 0.5\ E_{\text{cut}}$, and $E_{\text{cut}} = 4\epsilon$.

$U_{\text{c}}$ is introduced to mimic the confinement effect that chromosomes experience inside the cell due to their interaction with the nuclear envelope. It is defined as a spherical boundary whose radius $r_{\text{confine}}$ is chosen to maintain a given base pair density $\rho$. Each chromatin bead interacts with its nearest point on the boundary through a hard-core potential $U_{\text{c}}$.

$$U_{\text{c}} = \begin{cases} 4\epsilon\left[\left(\frac{\sigma}{r}\right)^{12} - \left(\frac{\sigma}{r}\right)^{6}\right] + \epsilon, r \leq \sigma 2^{1/6} \\ 0, r > \sigma 2^{1/6} \end{cases}$$

We initialize all the computer simulations with a chromatin configuration in which all beads are inside the boundary. Because of the hard-core potential, these beads will then remain inside the confinement during the entire simulation. The radius of the confinement is chosen to approximate the base pair density found in the human nucleus that encloses $\sim 6$ billion base pairs in a volume of $10\mu m$ in diameter. We therefore estimate the density as $\rho = \frac{6.176\times10^{9}\text{bp}}{\left(\frac{4}{3}\pi(5\mu\text{m})^{3}\right)} = 1.1795 \times 10^{7}\,\frac{bp}{\mu m^{3}}$, and set confinement size to be $0.79686\ \mu m = 26.562\ \sigma$ for the simulated chromosome segment of 25 Mb in length.

The two additional terms in Eq. [1] are introduced to model the genome organization starting from one-dimensional sequence features of the chromatin. In particular, $U_{\text{CS}}(\boldsymbol{r})$ quantifies chromatin state specific interaction energies between pairs of loci, and is defined as

$$U_{\text{CS}}(\boldsymbol{r}) = \sum_{IJ}\sum_{i\in I}\sum_{j\in J} \alpha^{IJ}(|j-i|)\, f(r_{ij}), \qquad [2]$$

where $I$ and $J$ indexes different chromatin states, and $i$ and $j$ runs over different genomic segments. This chromatin state-based potential is crucial for the formation of topologically associating domains (TADs) and the compartmentalization of chromatin domains.

$f(r_{ij})$ in Eq. [2] determines the probability to record a contact between a pair of genomic segments $i$ and $j$ separated by a distance $r_{ij}$, and is defined as follows

$$f(r) = \begin{cases} \frac{1}{2}\left[1 + \tanh\left(\sigma(r_c - r)\right)\right], \text{if } r \leq r_{\text{c}} \\ \frac{1}{2}\left(\frac{r_c}{r}\right)^{4}, \ \text{for } r > r_{\text{c}} \end{cases} \qquad [3]$$

where $r_c = 1.76$ and $\sigma = 3.72$. As shown in Fig. S8, compared to a simple hyperbolic tangent function used in previous studies [3, 4], the new expression decays to zero for large distances $r$ at a slower rate. This new form is motivated by the power law relationship between spatial distances and Hi-C contact probabilities observed in Ref. [5].

The prefactor $\alpha^{IJ}(|j-i|)$ in Eq. [2] measures the energetic cost for forming a contact between a pair of genomic loci $i$ and $j$. It depends explicitly on the chromatin states $I$ and $J$ for the two loci and also the genomic separation $|j-i|$ between them. As mentioned in the main text, the dependence of these contact energies on genomic separation is crucial to distinguish different mechanisms that lead to the formation of TADs at the intermediate scale, and the compartmentalization of chromatin domains separated far apart in sequence.

In its most general form, $\alpha^{IJ}(|j-i|)$ would require too many parameters to be parameterized robustly. For example, with 15 chromatin states and a chromosome of 4000 beads (20 Mb in sequence length), the parameter number would be $\frac{15(15+1)}{2} \times 4000 = 480000$. We therefore introduce the following strategy to reduce the number of parameters. First, we separate $\alpha^{IJ}(|j-i|)$ into two terms that include a mean field ideal potential $\alpha_{\text{ideal}}$ and a sequence specific potential $\alpha_{\text{seq}}$

$$\alpha^{IJ}(|j-i|) = \alpha_{\text{ideal}}(|j-i|) + \alpha^{IJ}_{\text{seq}}(|j-i|). \qquad [4]$$

We then approximate the two potential terms with different approaches. For the ideal potential, instead of defining its value at every sequence separation with a 5 kb increment, we use a coarsened grid with a spacing of 50kb and approximate all the potential in the same grid with a single value. For example,

$$\alpha_{\text{ideal}}(|j-i|) \equiv \alpha^{\text{coarse}}_{\text{ideal}}(10(s-1)+1)$$
$$\text{for } |j-i| \in [10(s-1)+1, 10s] \text{ and } s \in \left[1, \frac{N}{10}\right],$$

where $N$ is the total number of polymer beads. On top of the coarse grid, we add 40 more parameters to refine the ideal potential at a 5kb resolution for sequence separations $|j-i|$ that are less than 200 kb. The final expression for $\alpha_{\text{ideal}}(|j-i|)$ with $|j-i| \in [10(s-1)+1, 10s]$ can be summarized as

$$\alpha_{\text{ideal}}(|j-i|) = \begin{cases} \alpha^{\text{coarse}}_{\text{ideal}}(10(s-1)+1) + \alpha^{\text{fine}}_{\text{ideal}}(|j-i|), & s \le 40 \\ \alpha^{\text{coarse}}_{\text{ideal}}(10(s-1)+1), & s > 40. \end{cases}$$

A total of 440 parameters is thus used to define $\alpha_{\text{ideal}}(|j-i|)$.

For each pair of the chromatin states $I$ and $J$, we approximate the sequence specific potential with four piece-wise terms,

$$\alpha_{\text{seq}}^{IJ}(|j-i|) = \begin{cases} a_1^{IJ} + b_1^{IJ} \ln(|j-i|) + c_1^{IJ}|j-i|^{0.25}, & \text{for } |j-i| \le 1\text{Mb} \\ a_2^{IJ} + b_2^{IJ} \ln(|j-i|) + c_2^{IJ}|j-i|^{0.25}, & \text{for } 1\text{Mb} < |j-i| \le 2\text{Mb} \\ a_3^{IJ} + b_3^{IJ} \ln(|j-i|) + c_3^{IJ}|j-i|^{0.25}, & \text{for } 2\text{Mb} < |j-i| \le 5\text{Mb} \\ a_4^{IJ} + b_4^{IJ} \ln(|j-i|) + c_4^{IJ}|j-i|^{0.25}, & \text{for } 5\text{Mb} < |j-i| \le 20\text{Mb} \end{cases} \quad [5]$$

The functional form $a + b\ln(|j-i|) + c|j-i|^{0.25}$ is chosen to provide a good fit for the power-law decay $|j-i|^{\alpha}$ for most $\alpha \in [0,1]$ by varying the coefficients $a, b$ and $c$. Power-law decay is a reasonable approximation to the dependence of contact energies on sequence length separation. The four different sets of parameters make it possible to capture a possible change in $\alpha$ over the entire range of genomic separation studied. With the above definition, a total number of $\frac{15(15+1)}{2} \times 3 \times 4 = 1440$ parameters is needed for the sequence-specific term.

The last term in Eq. [1], $U_{\text{CTCF}}(\boldsymbol{r})$, is included to model the interaction between pairs of genomics loci $k$ and $l$ due to the formation of chromatin loops anchored by pairs of CCCTC-binding factor (CTCF), and is defined as

$$U_{\text{CTCF}}(\boldsymbol{r}) = \sum_{K,L} \sum_{K \le k < l \le L} \left[\alpha_1(1-\delta_{k,K})(1-\delta_{l,L}) + \alpha_2(\delta_{k,K} + \delta_{l,L})(1-\delta_{k,K}\delta_{l,L}) + \alpha_3\delta_{k,K}\delta_{l,L}\right] f(r_{kl}), \quad [6]$$

where $\delta$ is Kronecker delta function and $K$ ($L$) indexes over CTCF binding sites with 5' – 3'(3' – 5') orientation. The particular functional form for $U_{\text{CTCF}}(\boldsymbol{r})$ is motivated by the extrusion model [6-8]. For example, since binding of cohesin molecules to two strands of chromatin will bring them into spatial proximity, the effect of this binding can be approximated as an effective attraction between genomic loci. Since cohesin can extrude linearly along the chromosome within the region bound by a pair of convergent CTCF-binding sites, this effective interaction will also be limited to genomic loci enclosed by CTCF-binding sites (1st term). We used two additional parameters to model the interaction between CTCF-binding sites and the enclosed chromatin (2nd term), and interaction between CTCF-binding sites (3rd term). Since cohesin molecules have longer residence time near the CTCF molecules, the effective interaction between CTCF-binding sites and the chromatin may be stronger compared to chromatin-chromatin interaction, and therefore requires a separate potential term. We note that $U_{\text{CTCF}}(\boldsymbol{r})$ is only applied for convergent CTCF pairs that are separated by no more than 4 CTCF binding sites with 5' – 3' orientation or 4 CTCF binding sites with 3' – 5' orientation to mimic the finite processivity of cohesin molecules.

**Parameter optimization.** Though the chromatin energy function $U_{\text{Chrom}}(\boldsymbol{r})$ in Eq. [1] has clear physically meanings and is biologically justified, its expression can also be derived following the maximum entropy framework proposed in Refs. [3,4]. In particular, $U_{\text{Chrom}}(\boldsymbol{r})$ can be shown as the least biased functional form to reproduce the following set of constraints that include the average contact probabilities between a generic pair of genomic loci at various sequence separations, average contact probabilities between pairs of chromatin states $I$ and $J$ at various sequence separations, and average contact probabilities between pairs of CTCF binding sites $K$ and $L$ and between chromatin segments enclosed by convergent CTCF pairs

$$\begin{aligned}
\textstyle\sum_{i,j}\langle f(r_{ij})\delta_{|j-i|,s}\rangle &= \textstyle\sum_{i,j} f_{ij}^{\exp}\,\delta_{|j-i|,s} \\
\textstyle\sum_{i\in I}\sum_{j\in J}\langle f(r_{ij})\delta_{|j-i|,s}\rangle &= \textstyle\sum_{i\in I}\sum_{j\in J} f_{ij}^{\exp}\delta_{|j-i|,s} \\
\textstyle\sum_{K,L}\langle f(r_{KL})\rangle &= \textstyle\sum_{K,L} f_{KL}^{\exp} \\
\textstyle\sum_{K}\sum_{K<l<L}\langle f(r_{Kl})\rangle + \sum_{L}\sum_{K<k<L}\langle f(r_{kL})\rangle & \\
&= \textstyle\sum_{K}\sum_{K<l<L} f_{Kl}^{\exp} + \sum_{L}\sum_{K<k<L} f_{kL}^{\exp} \\
\textstyle\sum_{K,L}\sum_{K<k<l<L}\langle f(r_{kl})\rangle &= \textstyle\sum_{K,L}\sum_{K<k<l<L} f_{kl}^{\exp}
\end{aligned} \qquad [7]$$

$$\text{for } s = 1,\dots,N,\ \ I,J = 1\dots N_{\text{cs}},\ \ K = 1,\cdots,N_{\text{CTCF}}^{53}, \text{and } L = 1,\cdots,N_{\text{CTCF}}^{35}.$$

In the above equation, $\delta$ is the Kronecker delta function, $f_{ij}^{\exp}$ is the contact probability between the pair of genomic segments $i$ and $j$, $N$ is the number of polymer beads, $N_{\text{cs}}$ is the number of chromatin states, and $N_{\text{CTCF}}^{53}$ $N_{\text{CTCF}}^{35}$ are the number of CTCF binding sites oriented in the $5'-3'$ and $3'-5'$ directions respectively. The angular brackets $\langle\cdot\rangle$ represent ensemble averages over the Boltzmann distribution $e^{-\beta U_{\text{chrom}}(\boldsymbol{r})}$. Again, the summation over $K, L$ in the 3rd, 4th and 5th equation is only applied for convergent CTCF pairs that are separated by no more than 4 CTCF binding sites with 5' – 3' orientation or 4 CTCF binding sites with 3' – 5' orientation.

The first constraint in Eq. [7] will give rise to the ideal potential $\alpha_{\text{ideal}}(|j-i|)$ introduced in Eq. [4]. As mentioned above, $\alpha_{\text{ideal}}(|j-i|)$ is defined on a coarsened grid with a spacing of 50kb to reduce the number of parameters. Correspondingly, the constraints can be modified for this coarsened definition as

$$\sum_{i,j}\langle f(r_{ij})\Theta_{|j-i|,s}\rangle = \sum_{i,j} f_{ij}^{\exp}\,\Theta_{|j-i|,s}, \qquad \text{for } s = 1,\dots,\frac{N}{10}, \qquad [8]$$

where $\Theta_{|j-i|,s} = H\big(|j-i| - 10(s-1)\big) \times H(10s - |j-i|)$ and only has non-zero values in the region $|j-i| \in [10(s-1)+1, 10s]$. $H(x)$ is the Heaviside step function.

Similarly, we can adjust the second constraint in Eq. [7] for the approximate form of $\alpha_{\text{seq}}^{IJ}(|j-i|)$ defined in Eq. [5]. In particular, we can define the following constraints separately for each one of the four sequence ranges

$$\begin{aligned}
\sum_{i\in I}\sum_{j\in J}\langle f(r_{ij})\delta_{|j-i|,s}\rangle &= \sum_{i\in I}\sum_{j\in J} f_{ij}^{\exp}\delta_{|j-i|,s} \\
\sum_{i\in I}\sum_{j\in J}\langle f(r_{ij})\delta_{|j-i|,s}\ln|j-i|\rangle &= \sum_{i\in I}\sum_{j\in J} f_{ij}^{\exp}\delta_{|j-i|,s}\ln|j-i| \\
\sum_{i\in I}\sum_{j\in J}\langle f(r_{ij})\delta_{|j-i|,s}|j-i|^{0.25}\rangle &= \sum_{i\in I}\sum_{j\in J} f_{ij}^{\exp}\delta_{|j-i|,s}|j-i|^{0.25}
\end{aligned} \qquad [9]$$

The maximum entropy principle also suggests a simple optimization algorithm to derive the parameters in the potential. The strengths of contact energies, $\boldsymbol{\alpha}$, are determined by minimizing the objective function defined as:

$$\begin{aligned}\Gamma(\boldsymbol{\alpha}) = \ln\left(\frac{Z(\boldsymbol{\alpha})}{Z_0}\right) \\ + \beta\Bigg(\sum_s \alpha_{\text{ideal}}(s)\sum_{i,j}\delta_{|j-i|,s}f_{ij}^{\exp} + \sum_{I,J}\sum_s \alpha_{\text{residual}}^{IJ}(s)\sum_{i\in I}\sum_{j\in J}\delta_{|j-i|,s}\,f_{ij}^{\exp} \\ + \sum_{K,L}\sum_{K\le k<l\le L}\big[\alpha_1(1-\delta_{k,K})(1-\delta_{l,L}) + \alpha_2(\delta_{k,K}+\delta_{l,L})(1-\delta_{k,K}\delta_{l,L}) \\ + \alpha_3\delta_{k,K}\delta_{l,L}\big]f_{kl}^{\exp}\Bigg)\end{aligned} \quad [10]$$

where

$$Z(\boldsymbol{\alpha}) = \int d\boldsymbol{r}\; e^{-\beta U_{\text{Chrom}}(\boldsymbol{r})}$$
$$Z_o = \int d\boldsymbol{r}\; e^{-\beta U(\boldsymbol{r})}.$$

are the partition functions for $U_{\text{Chrom}}(\boldsymbol{r})$ and $U(\boldsymbol{r})$. For simplicity, $\Gamma(\boldsymbol{\alpha})$ is expressed in terms of the explicit constraints defined in Eq. [7]. Generalizing to the coarsened constraints defined in Eqs. [8] and [9] is straightforward. The objective function $\Gamma(\boldsymbol{\alpha})$ is a measurement of how much information theoretic entropy of the system is lost because of being constrained to the experimental input data. By maximizing $\Gamma(\boldsymbol{\alpha})$, the parameters in the potential $U_{\text{Chrom}}(\boldsymbol{r})$ can be found when $\frac{\partial\Gamma}{\partial\alpha} \equiv 0$. The same iterative algorithm outlined in Ref. [3] was used for parameter optimization. As mentioned in the main text, we used Hi-C experiments for segments of chromosomes 1, 10, 19, and 21 from GM12878 cells to determine the experimental constraints $f^{\exp}(r_{ij})$.

**Hi-C data analysis.** Experimental contact maps at 5kb resolution from Ref. [1] were downloaded using the Gene Expression Omnibus (GEO) accession number GSE63525. We used the combined contact matrices constructed from all read pairs that map to the genome with a MAPQ>=30. The raw matrices were then normalized with the KR method using the normalization vector provided in the same dataset. To convert the contact matrices into probabilities, we further divided each matrix element with the diagonal value $C_{ii} = 1035$ obtained from averaging over all chromosomes. With this probability conversion, all the genomic segments that are within in 5kb along the sequence will on average have a contact probability of 1. Since in the computational model, a 5kb segment has a diameter of $\sigma = 30$ nm, this probability conversion is equivalent of specifying the contact probability as 1 for genomic loci that are within a spatial distance of 30 nm. Such a probability definition is indeed consistent with the contact function $f(r)$ defined in Eq. [3] and plotted in Fig. S8.

**CTCF-binding sites from ChIP-Seq data.** We used the following protocol to process ChIP-Seq data and to extract genomic locations and orientations of CTCF binding sites.

Starting from the peak profile downloaded from ENCODE (see Extended data sheet for details), we identified the center of binding for each peak of both CTCF and Rad21. As both CTCF and cohesin molecules are found at the boundaries of most chromatin loops [9], we selected loop forming CTCF binding sites as those that have at least one Rad21 molecule located within 50bp of their genomic locations.

We then determined the orientation of each CTCF-binding site as follows. We first attempted to align the binding sites to the set of CTCF motifs compiled in Ref. [1, 10]. If the alignment succeeds and a motif is found within 100bp of the binding site, the orientation of the binding site was then assigned based on the DNA sequence of that motif. If no motif can be aligned, the orientation of the CTCF-binding site is determined using the genomic location of its binding center relative to that of the nearest binding center of Rad21. For example, we assign the orientation as 5' – 3' if the nearest Rad21 binding center is in the downstream of the CTCF binding site; otherwise, the orientation is assigned as 3' – 5'.

The above procedure will result in a list of oriented CTCF sites at single base resolution. From this list, we defined a 5kb-long bead in the computational model as a CTCF site if there is at least one CTCF binding site falls into the genomic region enclosed by that bead. If all the CTCF sites within the 5kb region have the 5' – 3' orientation, then the bead is assigned with the 5' – 3' orientation; similarly, if all the CTCF sites within the 5kb region have the 3' – 5' orientation, then the bead is assigned with the 3' – 5' orientation. If CTCF sites with both orientations are present, then the bead is assigned with dual orientation as well.

**Molecular dynamics simulation details.** All simulations were carried out using the molecular dynamics package LAMMPS [11] with reduced units $\sigma = 30\ nm$ and $\epsilon = k_B T$. Simulations were maintained at a constant temperature $T = 1.0$ via Langevin dynamics with a damping coefficient $\gamma = 0.5\tau$ and a time step of $dt = 0.012\tau$, where $\tau$ is the time unit.

Due to computational costs, we only simulated continuous genomic regions that are of 25Mb in length instead of whole chromosomes. These regions were mostly chosen from the q arms to avoid centromere regions that lack Hi-C data, and their genomic positions are provided in the Extended Data Sheet.

As aforementioned, the chromatin is confined in a spherical boundary during the simulation to mimic its interaction with the nuclear envelope. Such a confinement, however, also introduce boundary effects to the two end regions of the chromatin. Unlike the central regions, these two ends will experience more interaction with the boundary. To alleviate this boundary effect, we simulated a chromosome of 25Mb in length, but only used the middle 20Mb segment for analysis, and discarded the two end regions 0-2Mb and 22-25Mb.

To calculate the ensemble averages during the parameter optimization step, we carried out eight independent 20-million-time-step-long simulations for each one of the four chromosomes. The starting configurations for these simulations were chosen as the end configurations from the last iteration. All simulations in the first iteration started from a random polymer configuration confined in the spherical boundary.

For all the predictions, we carried out eight independent simulations, each of which lasted 40 million time steps.

**Contact enhancement metric for chromatin loops**. Chromatin loops are identified as pairs of genomic loci whose contact probabilities measured in Hi-C experiments are significantly higher than the local background signal. Based on this definition, we use the ratio of the average contact probability of the peak region over that of the local background region as a measure of loop prediction quality. For a loop to stand out from the background, this ratio should be significantly higher than that from a randomly selected pair of loci. An illustration of the definition of peak and local background region is provided in Fig. S3-1A. We define the peak and background regions as the areas enclosed by green and black squares respectively. The exact coordinates used to specify these regions relative to the genomic positions of the two loop anchors $(0, 0)$ are marked on the right panel of Fig. S3-1A. We define the average contact probabilities over all the pixels in the peak regions as $P_{\mathrm{p}}$, and the two background regions as $P_{\mathrm{b}}$. The contact enhancement of a chromatin loop is then calculated as the ratio of the contact probabilities $\frac{P_{\mathrm{p}}}{P_{\mathrm{b}}}$.

As a comparison, we also calculated the contact enhancements for a list of randomly selected genomic pairs. To generate these random pairs, we first selected their starting genomic positions from the simulated chromatin region with equal probability. The genomic separation for these random pairs were then determined based on a Gamma distribution obtained from a numerical fit to the length distribution of chromatin loops found in Hi-C contact maps, $P(l) = \frac{1}{b^a \Gamma(a)} l^{a-1} e^{-\frac{x}{b}}$, with $a = 1.69$ and $b = 175.19$ (see Figure S3-1B).

**Clustering analysis of simulated chromosome structures.** To investigate the spatial co-localization of different chromatin states, we performed the following clustering analysis using the algorithm introduced in Ref. [12]. Briefly, we identified the clusters as the set connected networks formed among genomic loci. Edges in these networks were defined between nearest neighbor loci as determined using Voronoi tessellation [13].

For simplicity, the clustering analysis was performed for five coarse chromatin types defined by grouping the 15 chromatin states introduced in Fig. 1: Active 1 (Tx3, Tx, TxEnhW, Tx5), Active 2 (TxEnh5, PromD1, PromU, EnhW1), Bivalent (PromBiv, EnhBiv, TxWk), Repressive (ReprPC, Het), and Inactive (ReprPCWk, Quies).

**Figures.**

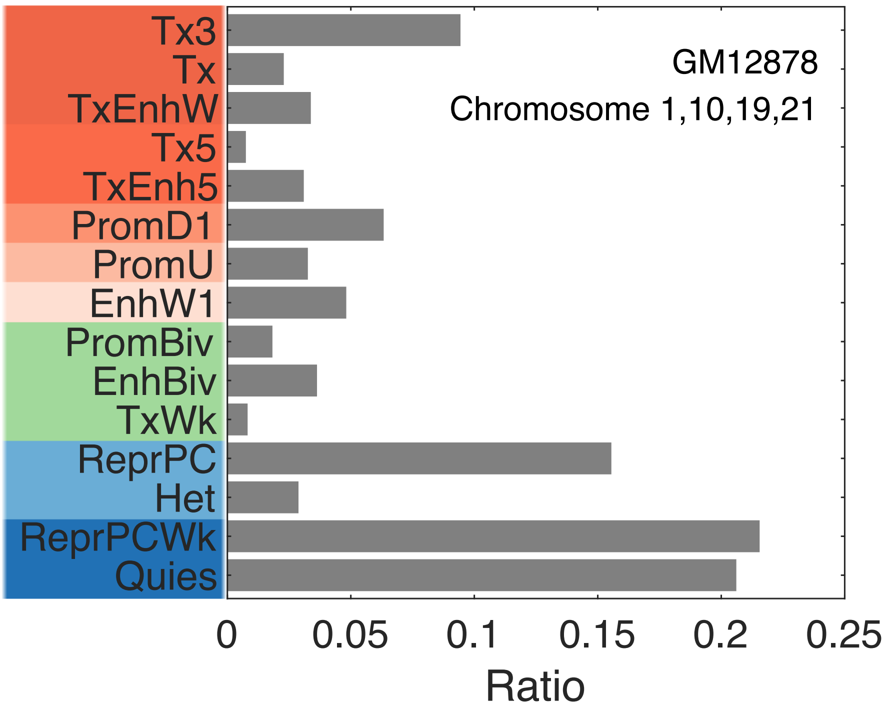


**Figure S1.** The coverage of various chromatin states in the selected segments of chromosomes 1, 10, 19, and 21 from GM12878 cells.

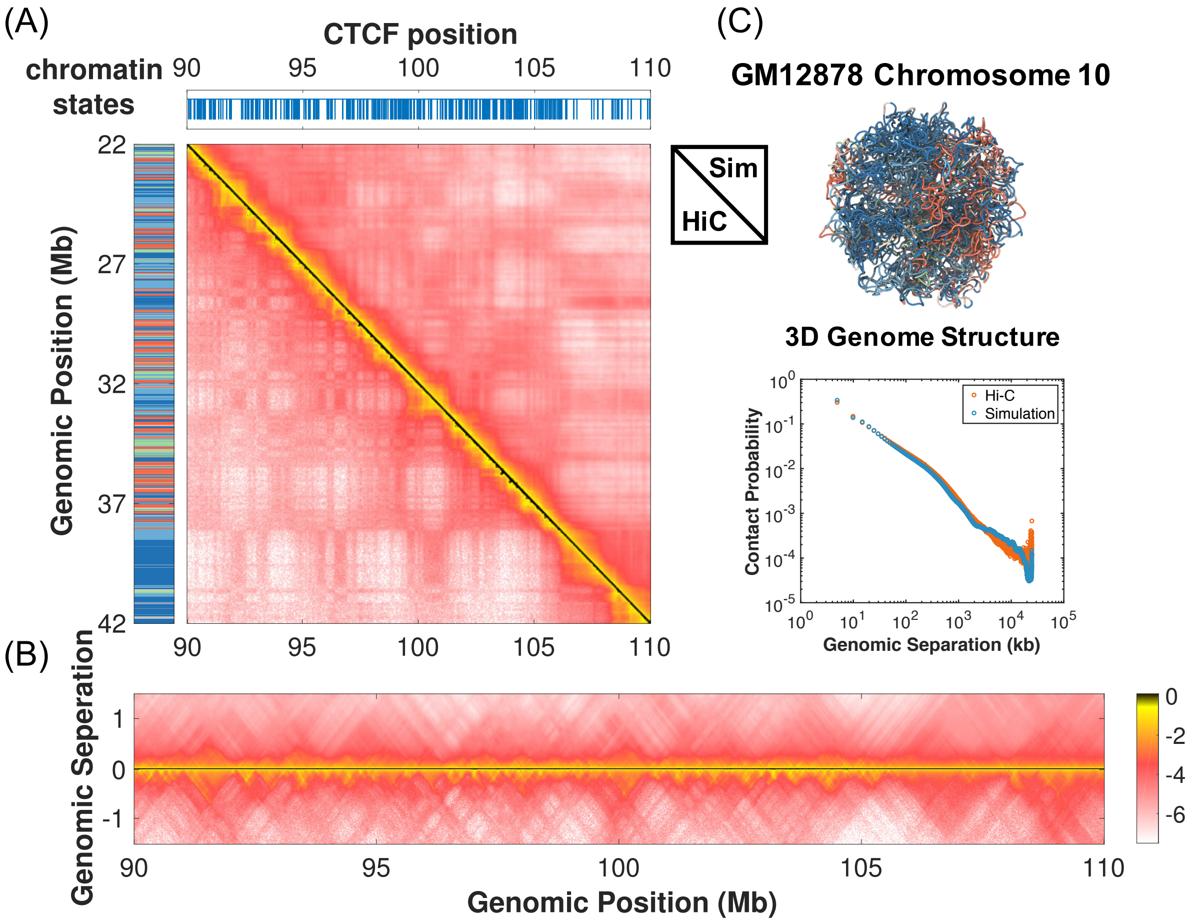


**Figure S2-1. Comparison between simulated and experimental contact probability maps for a 20 Mb segment of chromosome 10 from GM12878 cells.** (A) Results from simulation and the Hi-C experiment performed in Ref. [1] are shown in the upper and lower triangle respectively on a log scale. Also shown on the left and top panels are the sequence of chromatin states and the genomic positions of CTCF binding sites. A zoomed-in view of the contact maps along the diagonal region to highlight the formation of chromatin loops is shown in part (B). (C) A representative chromatin structure predicted by the computational model is drawn in a tube representation and colored by chromatin states. The average contact probability as a function of the genomic separation is shown below on a log-log scale for the simulated (blue) and experimental (red) contact maps respectively.

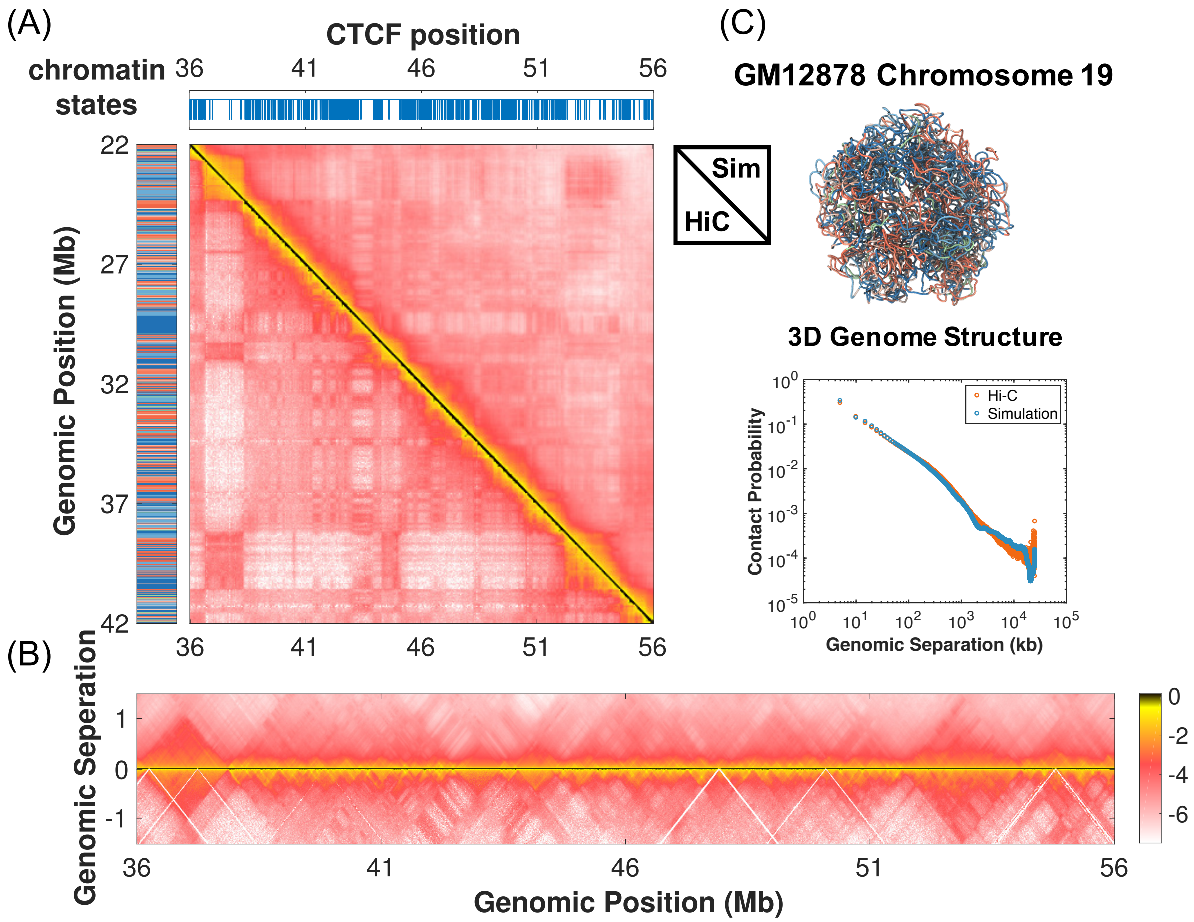


**Figure S2-2. Comparison between simulated and experimental contact probability maps for a 20 Mb segment of chromosome 19 from GM12878 cells.** (A) Results from simulation and the Hi-C experiment performed in Ref. [1] are shown in the upper and lower triangle respectively on a log scale. Also shown on the left and top panels are the sequence of chromatin states and the genomic positions of CTCF binding sites. A zoomed-in view of the contact maps along the diagonal region to highlight the formation of chromatin loops is shown in part (B). (C) A representative chromatin structure predicted by the computational model is drawn in a tube representation and colored by chromatin states. The average contact probability as a function of the genomic separation is shown below on a log-log scale for the simulated (blue) and experimental (red) contact maps respectively.

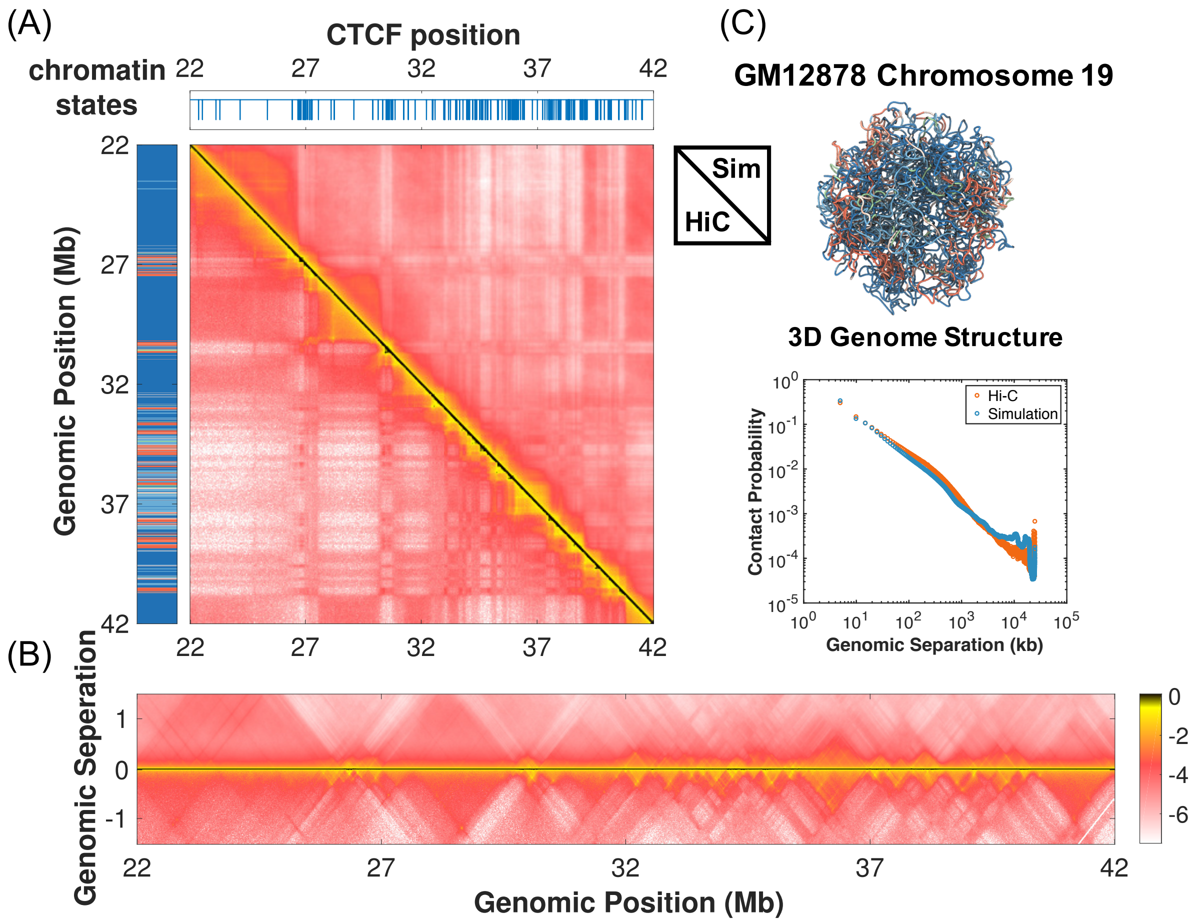


**Figure S2-3. Comparison between simulated and experimental contact probability maps for a 20 Mb segment of chromosome 21 from GM12878 cells.** (A) Results from simulation and the Hi-C experiment performed in Ref. [1] are shown in the upper and lower triangle respectively on a log scale. Also shown on the left and top panels are the sequence of chromatin states and the genomic positions of CTCF binding sites. A zoomed-in view of the contact maps along the diagonal region to highlight the formation of chromatin loops is shown in part (B). (C) A representative chromatin structure predicted by the computational model is drawn in a tube representation and colored by chromatin states. The average contact probability as a function of the genomic separation is shown below on a log-log scale for the simulated (blue) and experimental (red) contact maps respectively.

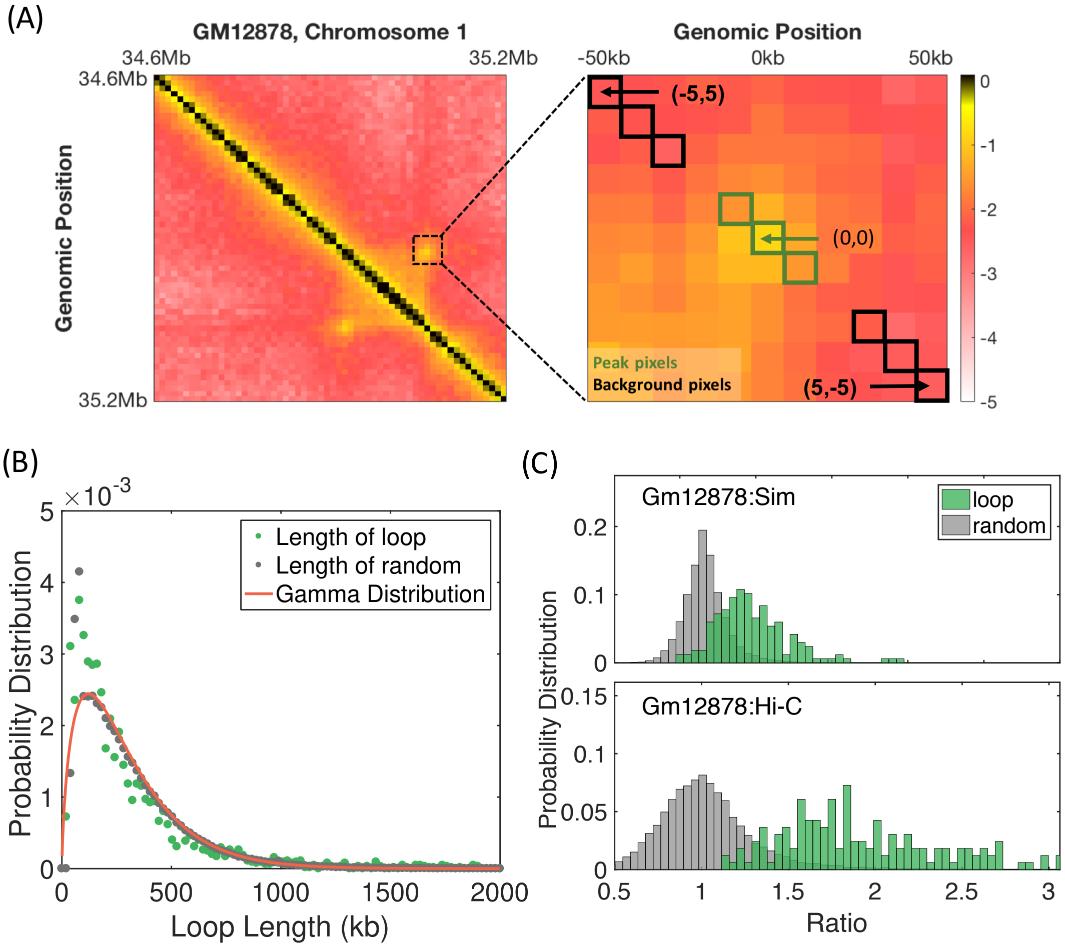


**Figure S3-1. Quantitative evaluation of the computational model's ability in predicting the formation of chromatin loops.** (A) Illustration for the definition of the contact enhancement metric introduced to evaluate the quality of chromatin loop prediction. An example contact probability map from Ref. [1] that highlights the formation of a chromatin loop between two genomic loci is shown on the left. A zoomed in view of the map is shown on the right to illustrate the definition of peak and background regions used to calculate the contact enhancement metric. See *SI Section: Contact enhancement metric for chromatin loops* for the definition of contact enhancement. (B) Probability distributions of the genomic separation for chromatin loops identified from Hi-C contact maps (green) and for randomly selected pairs (grey). The orange line is a numerical fit to the chromatin loop length distribution with a Gamma function. (C) Probability distributions of the contact enhancement for chromatin loops (green) and random pairs (grey) determined from the simulated (top) and the Hi-C contact map (bottom). Chromatin loops and contact maps from chromosomes 1, 10, 19 and 21 from GM12878 cells were used to calculate these distributions.

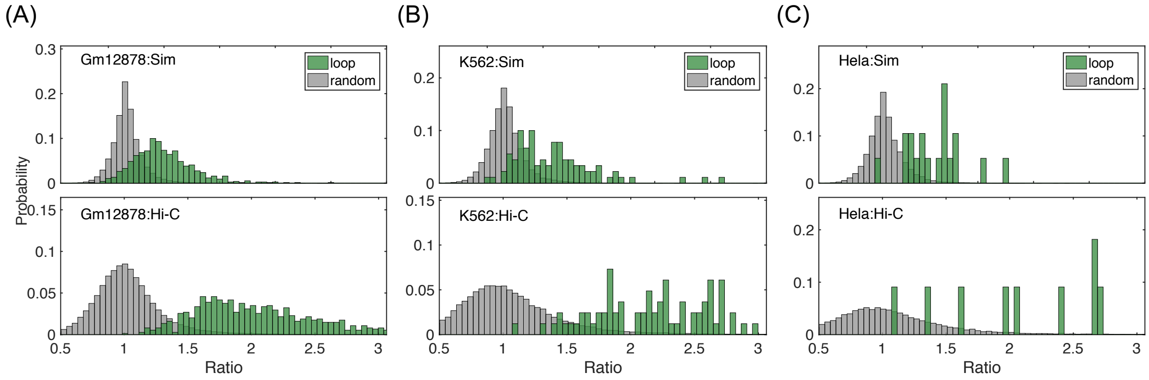


**Figure S3-2.** Probability distributions of the contact enhancement for chromatin loops (green) and random pairs (grey) determined from the simulated (top) and the Hi-C contact maps (bottom) for GM12878 (A), K562 (B) and Hela cells (C). Chromatin loops and contact maps from chromosomes 1 to 22 were used to calculate these distributions.

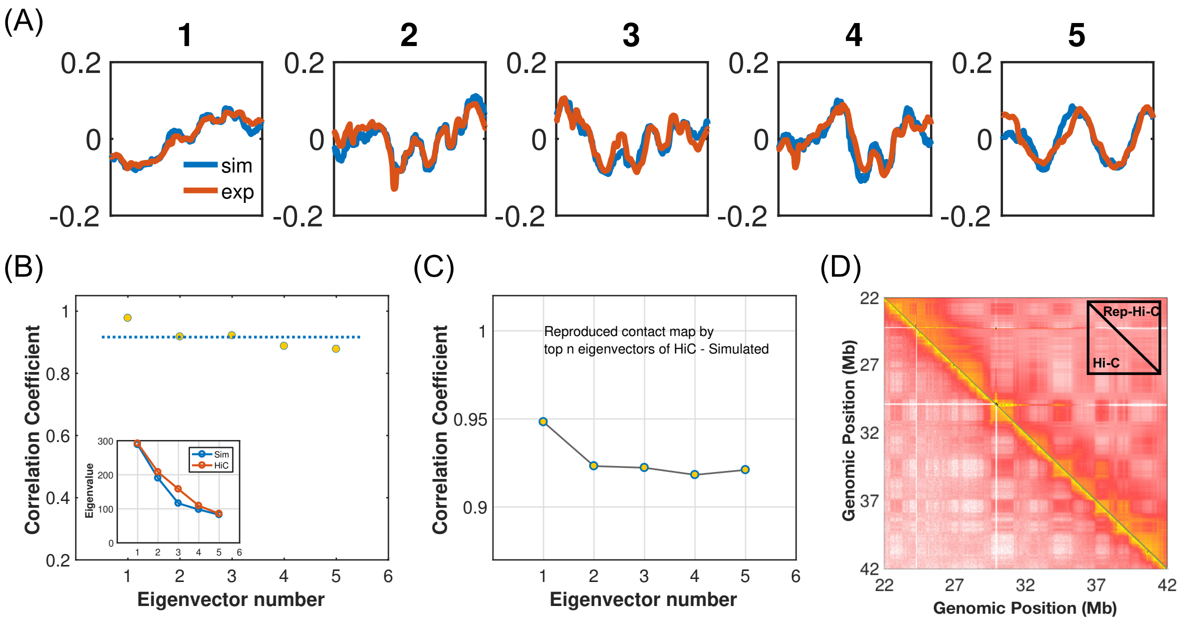


**Figure S4-1. Eigenvector analysis of the contact probability maps for chromosome 1 from GM12878 cells.** (A) Top five eigenvectors for simulated (blue) and experimental (orange) log contact matrices. (B) Correlation coefficients for the eigenvectors shown in (A). The average correlation coefficient (dotted line) is 0.917. The inset shows the comparison of the top five eigenvalues. (C) Correlation coefficients for the reproduced contact maps by the top $n$ eigenvectors of the experimental and simulated contact map as a function of $n$. (D) Contact map reconstructed using top five eigenvectors (top right) recapitulates the formation of TADs and compartments observed in the original map (bottom left).

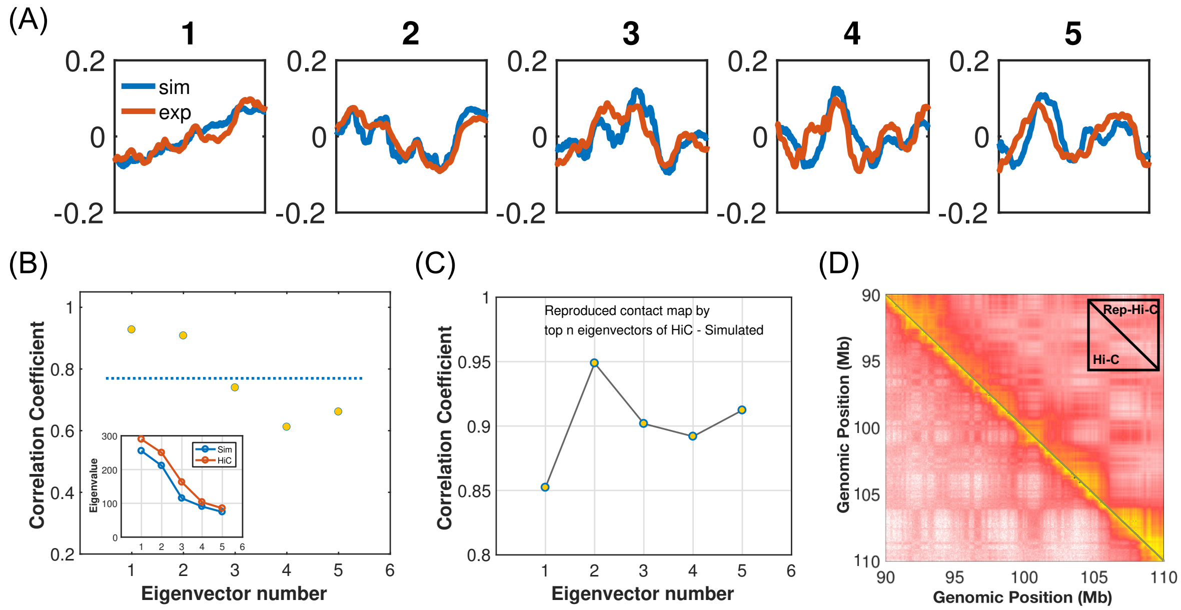


**Figure S4-2. Eigenvector analysis of the contact probability maps for chromosome 10 from GM12878 cells.** (A) Top five eigenvectors for simulated (blue) and experimental log contact matrices (orange). (B) Correlation coefficients for the eigenvectors shown in (A). The average correlation coefficient (dotted line) is 0.770. The inset shows the comparison of top five eigenvalues. (C) Correlation coefficients for the reproduced contact maps by the top $n$ eigenvectors of the experimental and simulated contact map as a function of $n$. (D) Contact map reconstructed using top five eigenvectors (top right) recapitulates the formation of TADs and compartments observed in the original map (bottom left).

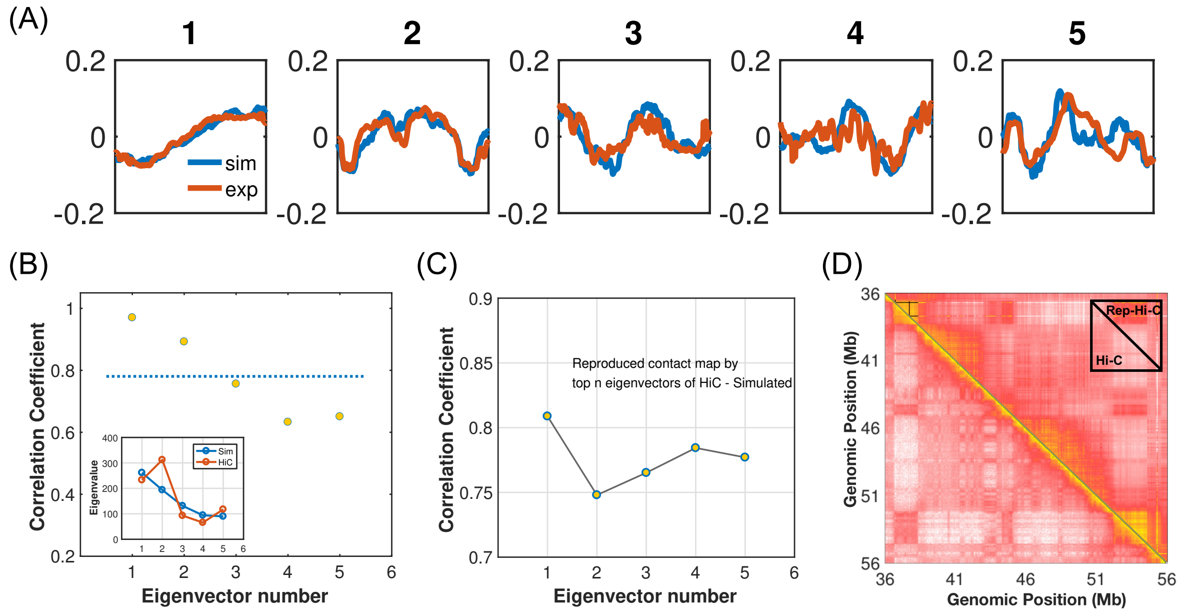


**Figure S4-3. Eigenvector analysis of the contact probability maps for chromosome 19 from GM12878 cells.** (A) Top five eigenvectors for simulated (blue) and experimental log contact matrices (orange). (B) Correlation coefficients for the eigenvectors shown in (A). The average correlation coefficient (dotted line) is 0.781. The inset shows the comparison of top five eigenvalues. (C) Correlation coefficients for the reproduced contact maps by the top $n$ eigenvectors of the experimental and simulated contact map as a function of $n$. (D) Contact map reconstructed using top five eigenvectors (top right) recapitulates the formation of TADs and compartments observed in the original map (bottom left).

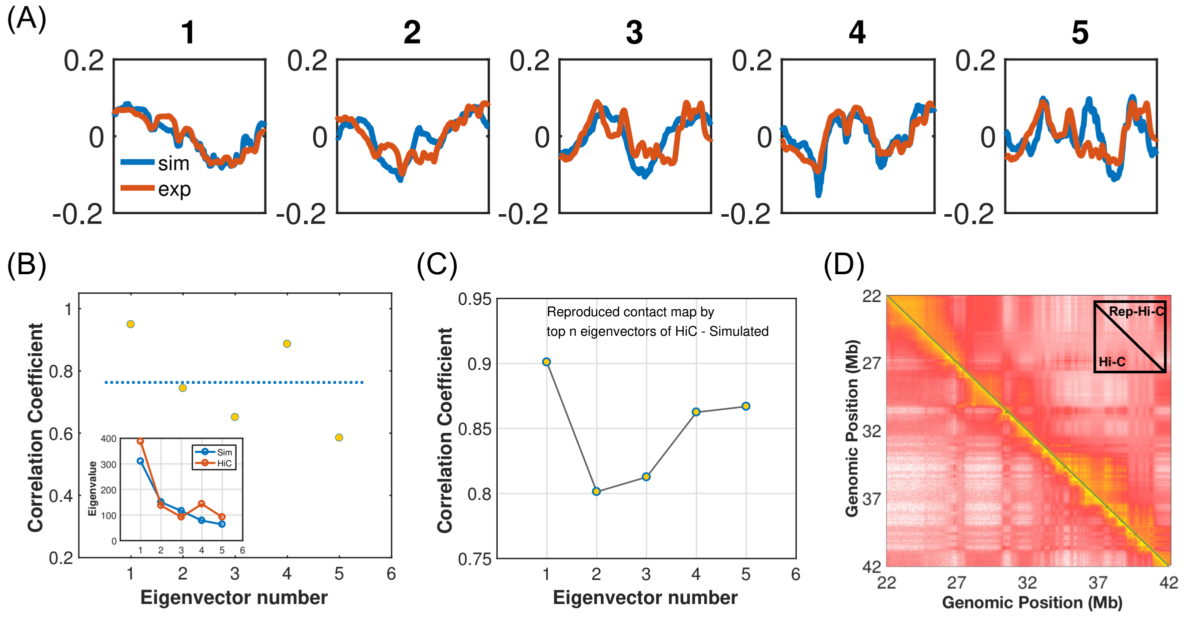


**Figure S4-4. Eigenvector analysis of the contact probability maps for chromosome 21 from GM12878 cells.** (A) Top five eigenvectors for simulated (blue) and experimental contact maps (orange). (B) Correlation coefficients for the eigenvectors shown in (A). The average correlation coefficient (dotted line) is 0.763. The inset shows the comparison of top five eigenvalues. (C) Correlation coefficients for the reproduced contact maps by the top $n$ eigenvectors of the experimental and simulated contact map as a function of $n$. (D) Contact map reconstructed using top five eigenvectors (top right) recapitulates the formation of TADs and compartments observed in the original map (bottom left).

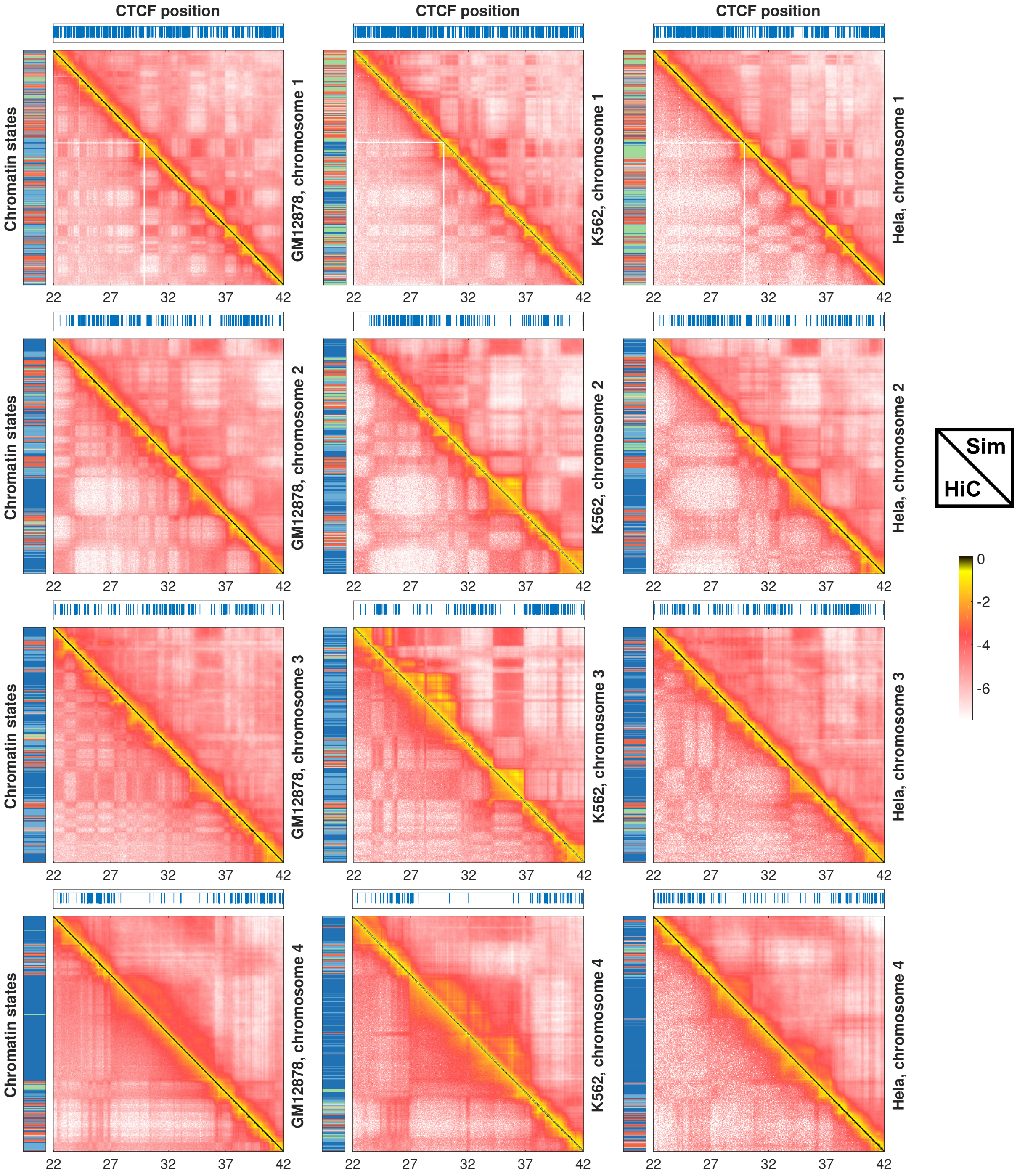


**Figure S5-1. Comparison between experimental (bottom left) and simulated (top right) contact maps for chromosomes 1-4 from GM12878 (left), K562 (middle) and Hela (right) cells.** Also shown on the left and top panels are the sequence of chromatin states and the genomic positions of CTCF binding sites.

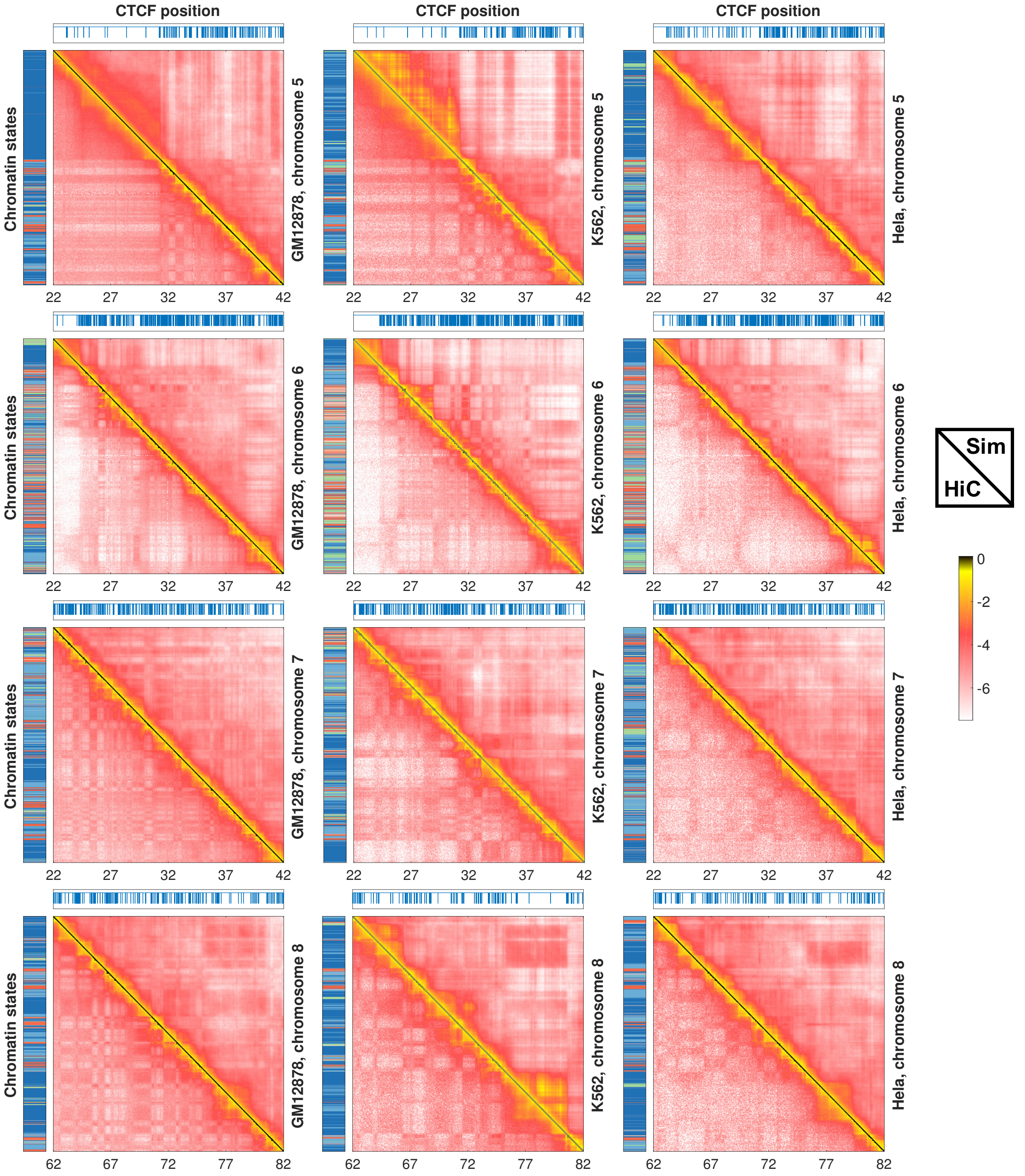


**Figure S5-2. Comparison between experimental (bottom left) and simulated (top right) contact maps for chromosomes 5-8 from GM12878 (left), K562 (middle) and Hela (right) cells.** Also shown on the left and top panels are the sequence of chromatin states and the genomic positions of CTCF binding sites.

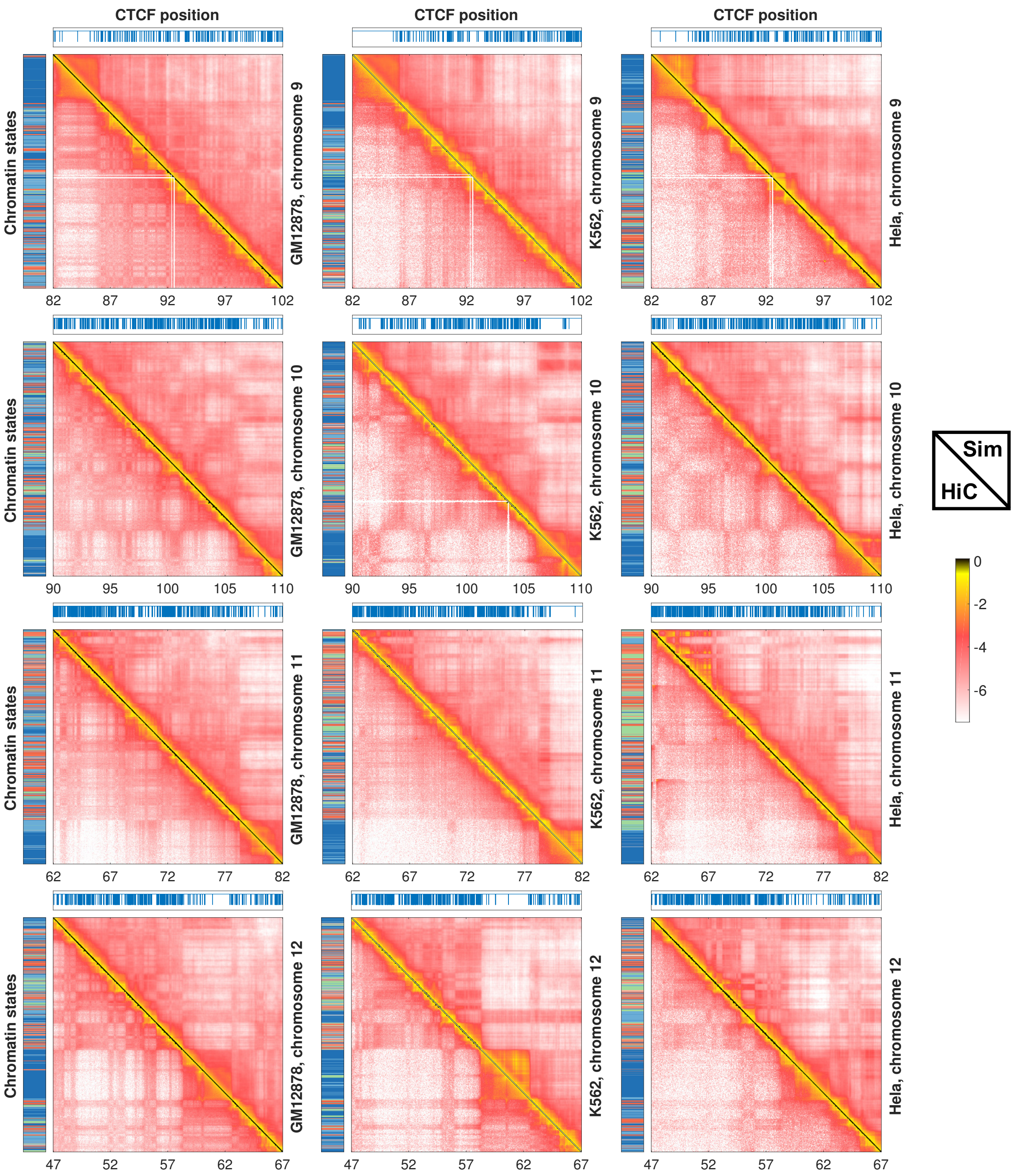


**Figure S5-3. Comparison between experimental (bottom left) and simulated (top right) contact maps for chromosomes 9-12 from GM12878 (left), K562 (middle) and Hela (right) cells.** Also shown on the left and top panels are the sequence of chromatin states and the genomic positions of CTCF binding sites.

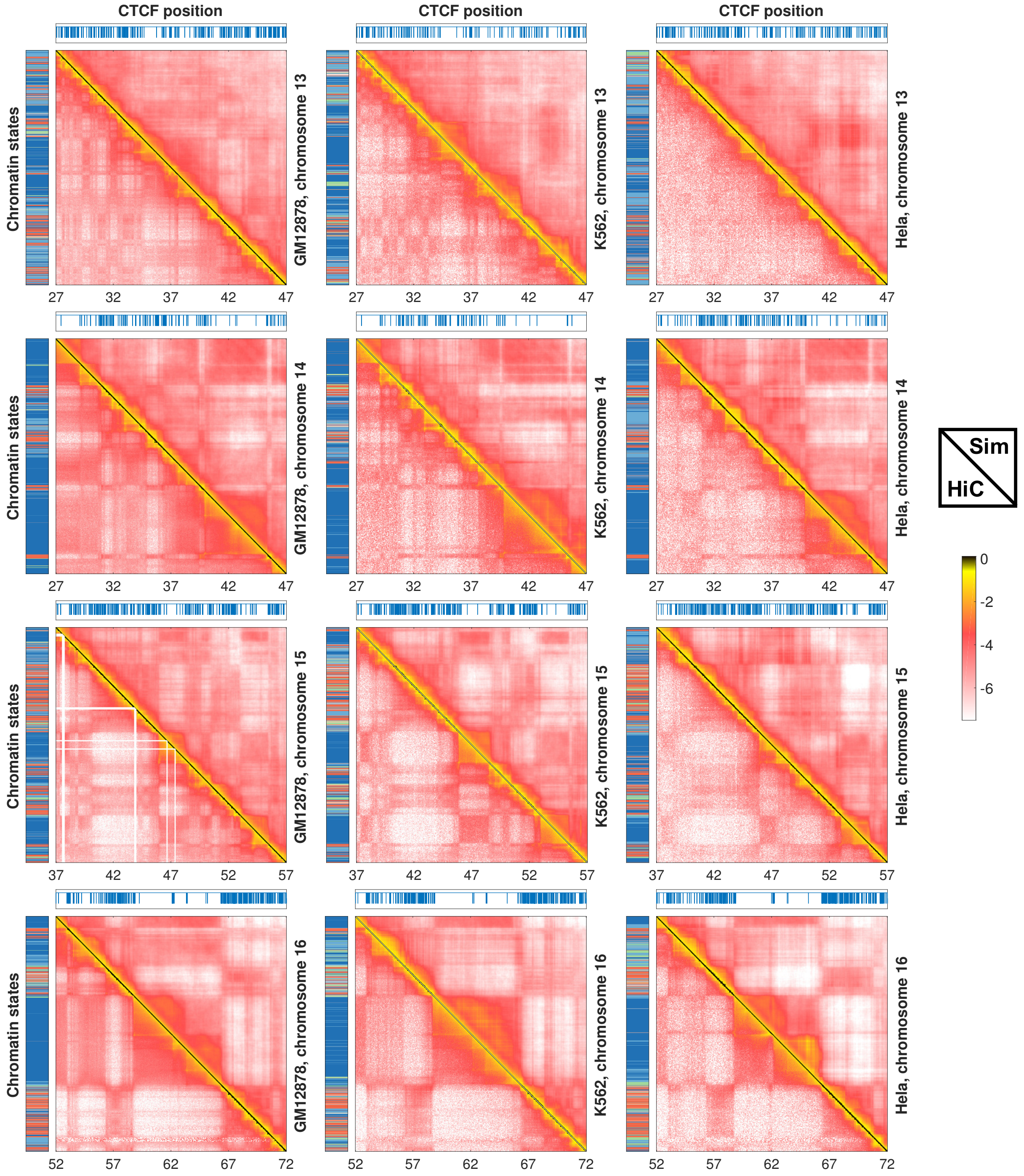


**Figure S5-4. Comparison between experimental (bottom left) and simulated (top right) contact maps for chromosomes 13-16 from GM12878 (left), K562 (middle) and Hela (right) cells.** Also shown on the left and top panels are the sequence of chromatin states and the genomic positions of CTCF binding sites.

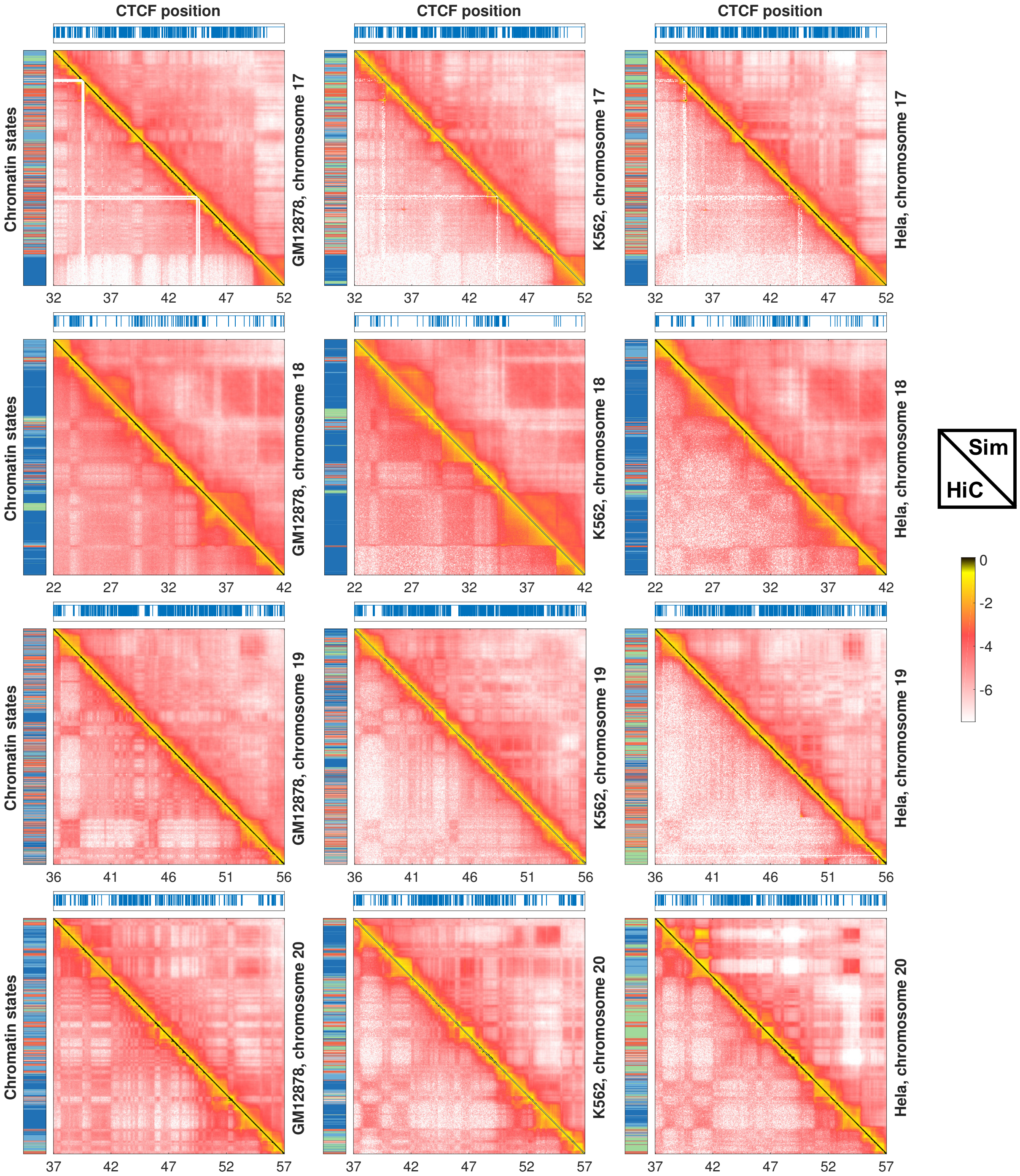


**Figure S5-5. Comparison between experimental (bottom left) and simulated (top right) contact maps for chromosomes 17-20 from GM12878 (left), K562 (middle) and Hela (right) cells.** Also shown on the left and top panels are the sequence of chromatin states and the genomic positions of CTCF binding sites.

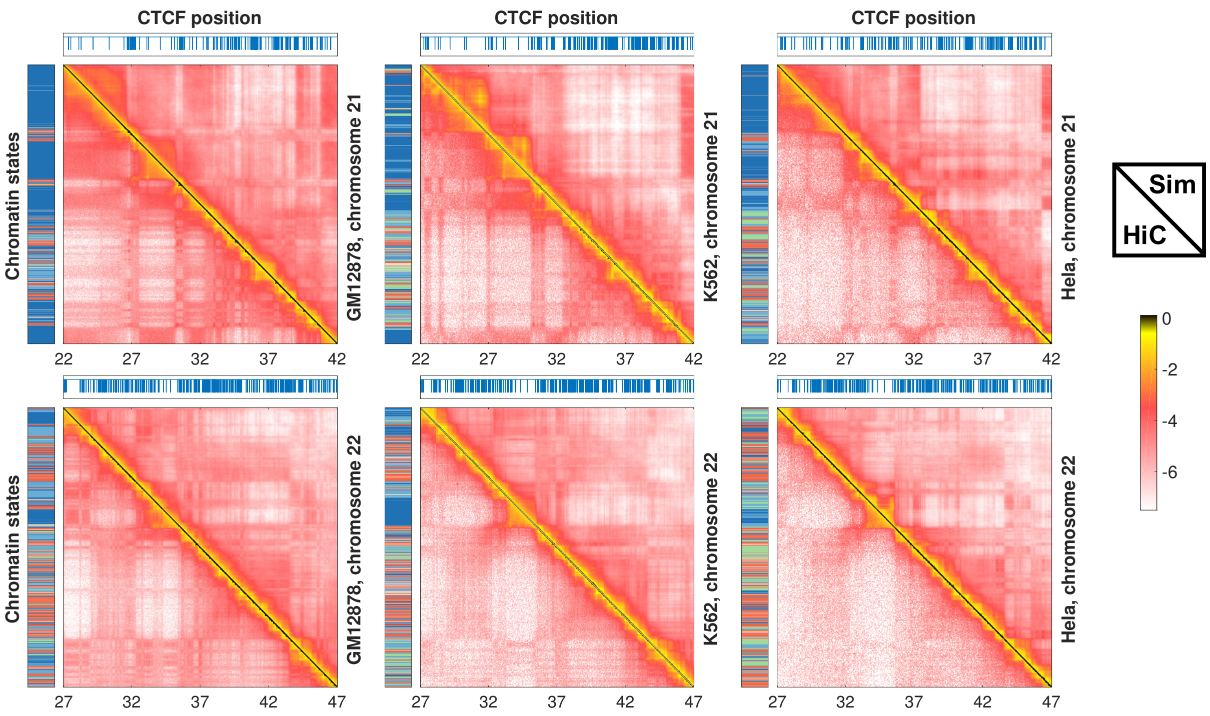


**Figure S5-6. Comparison between experimental (bottom left) and simulated (top right) contact maps for chromosomes 21-22 from GM12878 (left), K562 (middle) and Hela (right) cells.** Also shown on the left and top panels are the sequence of chromatin states and the genomic positions of CTCF binding sites.

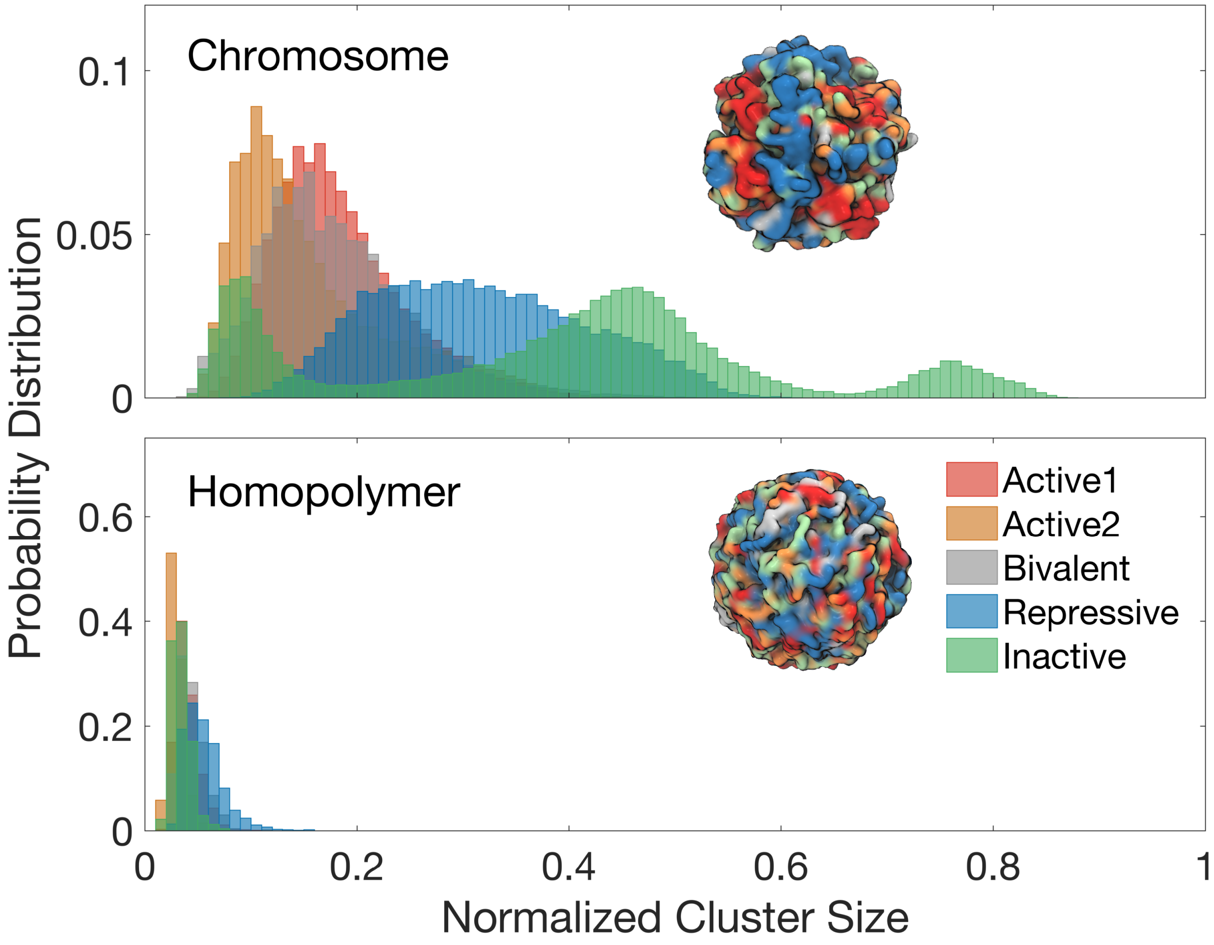


**Figure S6. Clustering analysis of simulated chromosome structures.** The probability distribution of the cluster size for different chromatin types calculated using the structural ensemble for chromosomes 1, 10, 19, 21 from GM12878 cells and for a homopolymer of the same length (*Bottom*). The largest two connected networks were used to determine the cluster sizes. See *SI Section: Clustering analysis of simulated chromosome structures* for details of the clustering algorithm. Representative chromosome and homopolymer structures colored by chromatin types are shown in the insets with a surface representation. Comparing the results shown in the top and bottom panel, it is clear that there is a tendency for genomic loci of the same chromatin type to co-localize spatially.

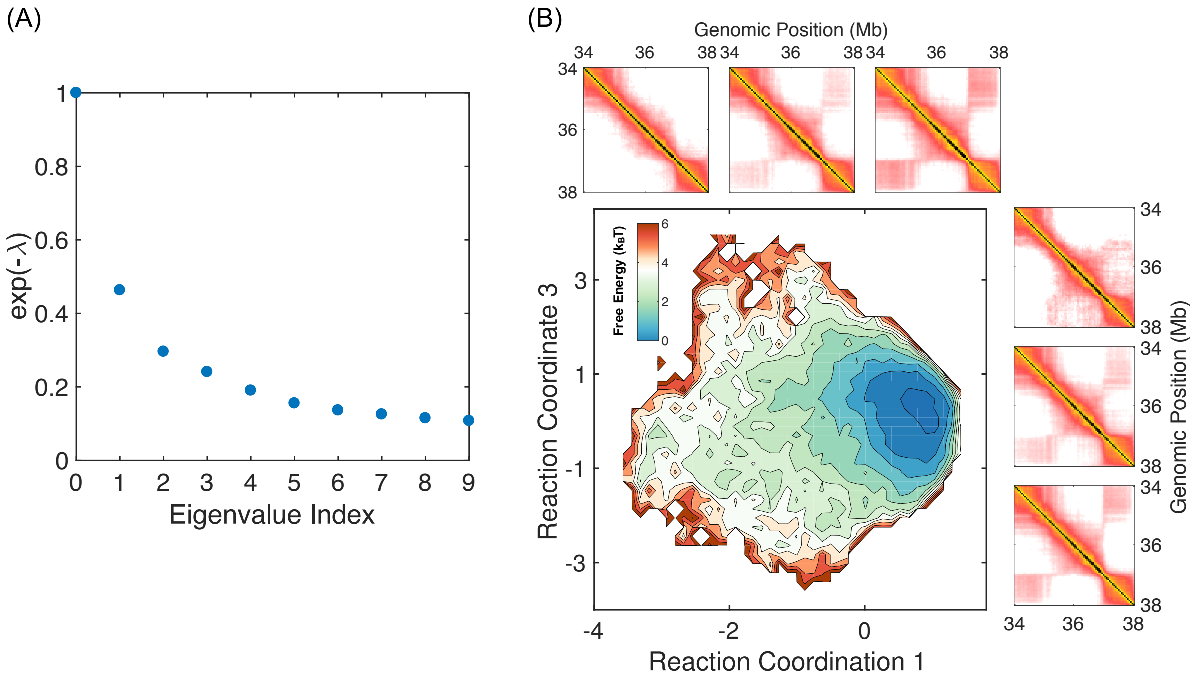


**Figure S7.** (A) Negative exponential of the eigenvalues of the transition matrix calculated in the diffusion map analysis of the genomic region chr1:34-38Mb from GM12878 cells. The eigenvectors corresponding to the second ($\lambda_1$) and third ($\lambda_2$) eigenvalues are selected to serve as reaction coordinates 1 and 2 respectively. (B) Free energy profile of TAD conformations projected onto eigenfunctions that correspond to the 1$^{st}$ and the 3$^{rd}$ eigenvalues. The (Left) and (Top) panels illustrate the change in contact maps along the two coordinates. The three contact maps for reaction coordinate 1 were identical to those shown in Figure 6, and the three regions used to calculated the contact maps for reaction coordinate 3 are $[-3.0, 0), [0, 1.5)$, and, $[1.5, 3.0)$.

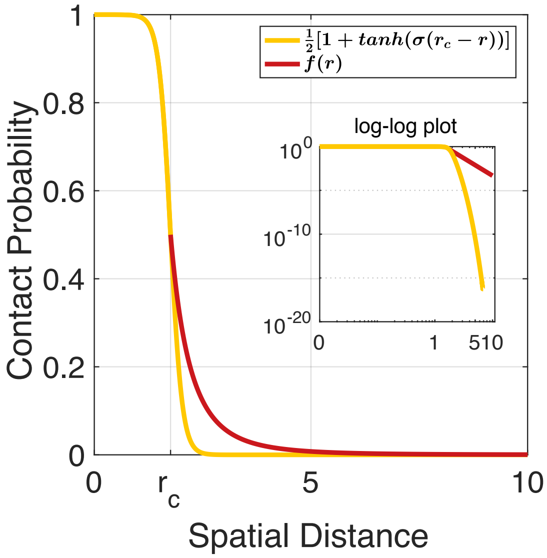


**Figure S8.** The probability of contact formation as measured by the function $f(r)$ defined in Eq. [3] and by a simple switching function. The same plot is shown in the inset on a log-log scale.

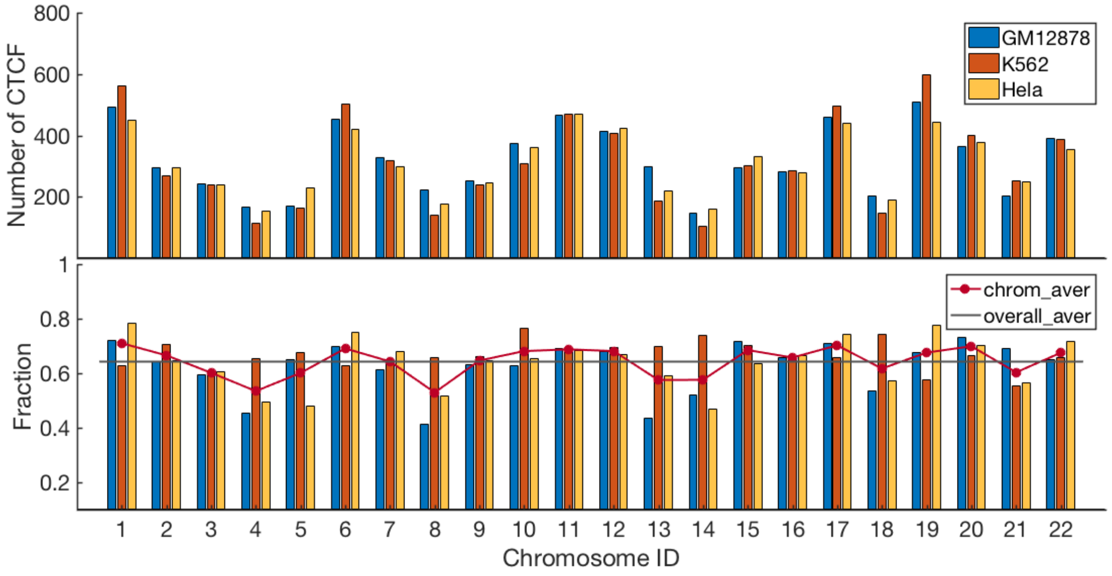


**Figure S9. Statistics of CTCF-binding sites for different cell types.** Number of CTCF-binding sites for chromosomes from GM12878 (blue), K562 (orange), and Hela (yellow) cells. (*Bottom*) The fraction of conserved CTCF-binding sites across the three cell types. The red dots are the average fractions over the three cell types for different chromosomes, and the grey line indicates the average over all chromosomes.
